# The geometry of domain-general performance monitoring in the human medial frontal cortex

**DOI:** 10.1101/2021.07.08.451594

**Authors:** Zhongzheng Fu, Danielle Beam, Jeffrey M. Chung, Chrystal M. Reed, Adam N. Mamelak, Ralph Adolphs, Ueli Rutishauser

**Affiliations:** Department of Neurosurgery, Cedars-Sinai Medical Center, Los Angeles, CA, USA; Division of Humanities and Social Sciences, California Institute of Technology, Pasadena, CA, USA; Department of Neurology, Cedars-Sinai Medical Center, Los Angeles, CA, USA; Division of Biology and Bioengineering, California Institute of Technology, Pasadena, CA, USA; Center for Neural Science and Medicine, Department of Biomedical Sciences, Cedars-Sinai Medical Center, Los Angeles, CA, USA

## Abstract

Controlling behavior to flexibly achieve desired goals depends on the ability to monitor one’s own performance. It is unknown how performance monitoring can be both flexible to support different tasks and specialized to perform well on each. We recorded single neurons in the human medial frontal cortex while subjects performed two tasks that involve three types of cognitive conflict. Neurons encoding predicted conflict, conflict, and error in one or both tasks were intermixed, forming a representational geometry that simultaneously allowed task specialization and generalization. Neurons encoding conflict retrospectively served to update internal estimates of control demand. Population representations of conflict were compositional. These findings reveal how representations of evaluative signals can be both abstract and task-specific and suggest a neuronal mechanism for estimating control demand.

## Introduction

Humans can rapidly learn to perform novel tasks given only abstract rules, even if task requirements differ drastically. To achieve this, cognitive control needs to coordinate processes across a diverse array of perceptual, motor and memory domains and at different levels of abstraction over sensorimotor representations (*1–3*). A key component of cognitive control is performance monitoring, which enables us to constantly evaluate whether we have made an error, experienced conflict, and responded fast or slow (*4, 5*). It provides task-specific information about which processes cause an error or a slow response so that they can be selectively guided (*6–15*), thereby solving the problem of “credit assignment” (*13, 16, 17*). At the same time, performance monitoring needs to be flexible and domain-general to enable cognitive control for novel tasks (*18*), to inform abstract strategies (e.g., “win-stay, lose-switch”, exploration versus exploitation (*19, 20*)), and to initiate global adaptation in motor (*21–23*), arousal (*24–26*) and emotional (*27–29*) states. As an example, errors and conflicts can have entirely different causes in different tasks (task-specific) but all signify failure or difficulty to fulfill an intended abstract goal (task-general); performance monitoring should satisfy both requirements. Enabled by its broad connectivity with the lateral prefrontal cortex (LPFC), thalamic and brainstem structures (*30, 31*), the medial frontal cortex (MFC) serves a central role in evaluating one’s own performance and decisions (*4, 10, 13, 17, 20, 25, 32–43*). However, little is known about how neural representation in the MFC can support both domain-specific and domain-general adaptations. Understanding the underlying mechanisms requires recording from the same neurons across multiple tasks, which has not been done.

Specialization and generalization place different constraints on neural representations (*44, 45*). Specialization demands separation of encoded task parameters, which can be fulfilled by increasing the dimensionality of neural representation (*46, 47*). By contrast, generalization involves abstracting away details specific to performing a single task, which requires reducing the representational dimensionality (*3, 46*). Theoretical work shows that the geometry of population activity can be configured to accommodate both of these seemingly conflicting demands (*44*), provided that the constituent single neurons multiplex task parameters non-linearly (*48, 49*). While recent experimental work has shown that neuronal population activity is indeed organized this way in the frontal cortex and hippocampus in macaques (*44*) and humans (*50*) during decision making tasks, it remains unknown whether this framework is applicable to the important topic of cognitive control. Here, we examine the hypothesis that neuronal populations in the human MFC represent performance monitoring in such a format, thereby enabling downstream processes for both domain-specific and domain-general control.

A key aspect of behavioral control is learning about the identity and intensity of control needed to correctly perform a task and deploy control proactively based on such estimates. Doing so requires integrating performance outcomes over multiple trials (*13*). BOLD-fMRI studies localize signals related to control demand estimation to the insular-frontostriatal network (*51*). However, no single-neuron substrate of these signals is known. Also, it is unknown how these slowly varying state-like signals are updated after a trial and whether the underlying substrate is domain-general or task-specific. We constructed a normative Bayesian model that captures this learning process as iterative estimation of the probability of encountering a specific type of conflict trial. A key prediction is that conflict is not only represented *ex ante* before an action is produced, but also *ex post* as an outcome signal. Our model also predicts that estimated conflict probability is slowly updated and relatively stable across adjacent trials. We tested these predictions by tracking single neuron activity within the MFC while patients performed two well-studied conflict tasks: the color-word Stroop task and the Multi-Source Interference Task (MSIT, (*52*)).

## Results

### Task and behavior

Subjects (see **Table S1**) performed the MSIT and the Stroop task (**Fig. 1A** and **Methods**; some patients only performed one task due to time constraints, **Table S1**). Conflict and errors arose from different sources in these two tasks: competition between a prepotency of reading over color naming in the Stroop task, and competition between the target response and either the spatial location of target (“Simon effect”, denoted by “fl”) or flanking stimuli (“Flanker effect”, denoted by “si”), or both (“sf”) in MSIT. In the MSIT, we refer to trials with or without a Simon Effect conflict as “Simon” and “non-Simon” trials, respectively (and similarly for Flanker trials). Stimulus sequences were randomized, with each type of trial occurring with a fixed probability (Stroop: 33% were conflict trials; MSIT: 15%, 15% and 30% had si, fl, and sf types of conflict, respectively; see **Methods** for details).

**Figure 1.**
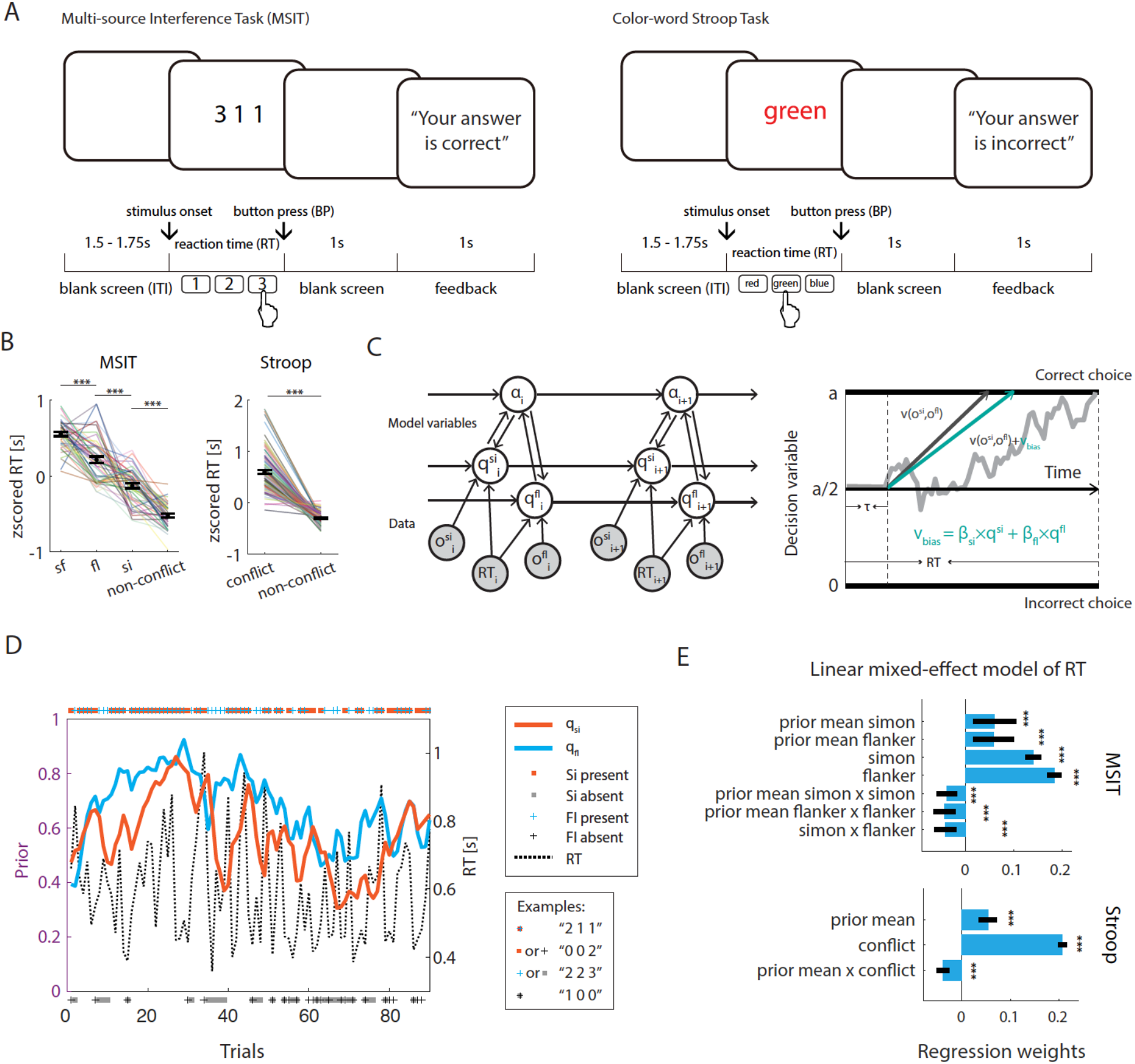
Tasks, Bayesian conflict learning model, reaction time analyses. (A) Tasks. (B) RTs were significantly prolonged by conflict in MSIT (left, N=41 sessions) and Stroop (right, N=82 sessions). (C) The updating process (left) and the decision process modelled as a drift diffusion process (right). Shown is the MSIT model, which has the five variables the flexible learning rate (a), predicted Simon conflict (q_si_), predicted Flanker conflict (q_fl_), observed Simon conflict (o_si_), observed Flanker conflict (o_fl_), and RT. Observables (trial congruency, RT, and outcome) are shown in gray, internal variables in white. Arrows indicate information flow. Horizontal and top-down edges represent the estimation of flexible learning rate, predicted conflict before stimulus onset, and the bottom-up edges represent using the observed conflict labels and RT to update latent variables for the next trial. (D) Estimated mean of the prior for Simon probability (orange) and Flanker probability (blue) from an example MSIT session. Markers placed on the top indicated the type of conflict present. (E) Regression analyses of RT using linear mixed-effect models. Blue bars show regression coefficients; black bars show confidence intervals. Conflict prior positively predicted RT in MSIT (top) and Stroop (bottom). *p < 0.05, ** p < 0.01, *** p < 0.001, n.s., not significant (p > 0.05).

Subjects performed well, with an error rate of 6.3±5% and 6.0±5% in the Stroop and MSIT tasks, respectively. Reaction times (RT) on correct trials were significantly prolonged in the presence of conflicts (**Fig. 1B**, see figure for statistics). Participants’ sequential performance (RT and accuracy) were modeled with a Bayesian conflict learning framework, building on existing models (*51, 53, 54*). Our model (**Fig. 1C)** assumes that participants iteratively estimated how likely they were to encounter a certain type of conflict (conflict probability) by updating their prior estimate with the new evidence received (experienced conflict and RT) on each trial. Since trial sequences were randomized, subjects could only estimate the probability that an upcoming trial would involve a conflict, an experimental parameter that was fixed but unknown to the subject a priori. RT generation was modeled as a drift-diffusion process (DDM), with drift rates depending both on the type of conflict present and the current estimate of the priors. In the following analyses, we refer to the means of the prior and posterior distributions as conflict “prior” (before stimulus onset) or “posterior” (after action completion). We fit the combined model, which encompasses both the Bayesian component and the DDM, to the behavior (RT, conflict) of our subjects using expectation-maximization (see **Methods**; See **Fig. S2** for estimated DDM hyperparameter values). To illustrate this process, **Fig. 1D** shows the measured behavior and the model-derived prior estimates of an example MSIT session.

We validated the behavioral relevance of the Bayesian model by examining how well the model-derived regressors predicted RT and errors. First, in addition to current trial conflict, the model-estimated conflict prior had a significantly positive main effect on RT in both tasks (i.e., an increase in RT; **Fig. 1E;** χ^2^(1) = 6.75, p = 0.009 for Simon and χ^2^(1) = 6.79, p = 0.009 for Flanker in MSIT; χ^2^(1) = 28.1, p < 0.001 for Stroop; likelihood ratio test). The extent to which RT varied with the conflict prior depended on the type of conflict (in the case of MSIT, Simon and Flanker separately), as indicated by a significant negative interaction term (χ^2^(1) = 12.94 for Simon and χ^2^(1) = 14.2 for Flanker in MSIT; χ^2^(1) = 33.3 for Stroop. p < 0.001 for all conflict types; likelihood ratio test). This relationship between conflict prior and RT remained statistically significant when trial ID was added as nuisance variable (**Fig. S1A**), and when the conflict prior was estimated without RT tuning (**Fig. S1B**; see **Methods** for details). Our model yielded qualitatively similar results when applied to data from a separate group of healthy control subjects (**Fig. S1D**; the same RT-tuned models were used). Also, model-derived conflict prior was systematically related to errors: when conflict was estimated to be likely, subjects were *less* likely to commit an error on this trial, suggesting that more control was engaged (**Fig. S1C**; for MSIT we only considered “sf” trials where most errors occurred; significant main effect χ^2^(1) = 6.81, p = 0.009 for MSIT; significant interaction with non-significant main effect χ^2^(1) = 18.59, p < 0.001 for Stroop; likelihood ratio test). Collectively, these behavioral data from two tasks demonstrate that the iteratively estimated conflict probabilities were correlated with prolonged RT and reduced error likelihood, consistent with prospective engagement of control.

We further validated our model by comparing it with alternative models. First, our model explained significantly more variance in RT than a model that only has access to whether the previous trial was a conflict trial or not (**Table S2**, right). This finding, consistent with prior work (*51*), demonstrates that our participants incorporated conflict information from multiple previous trials. Second, we considered two alternative classes of models with more free parameters: a reinforcement learning model that performed trial-by-trial updating using a constant learning rate (free parameter) and a model that did not update trial-by-trial (free parameter is the constant conflict probability; “no updating” in **Table S3**). Parameters of these alternative models were optimized by maximum likelihood (MLE), which required access to all data. Our model performed significantly better than either class of alternative models (**Tables S2** and **S3** for a summary of model comparisons). RT tuning significantly improved our model both in terms of explaining RT (**Table S2**) and trial congruency (**Table S3**). Since the RT-tuned Bayesian models explained the most behavioral variance of all models considered, we used them for all neural analyses.

### Neuronal correlates of performance monitoring signals

We collected single-neuron recordings from two regions within the MFC (**Fig. 2A**): the dorsal anterior cingulate cortex (dACC) and the pre-supplementary motor area (pre-SMA). We isolated in total 1431 putative single neurons (Stroop: 593 in the dACC and 607 in the pre-SMA across 32 participants (10 females); MSIT: 326 in the dACC and 412 in the pre-SMA in 12 participants (6 females)).

**Figure 2.**
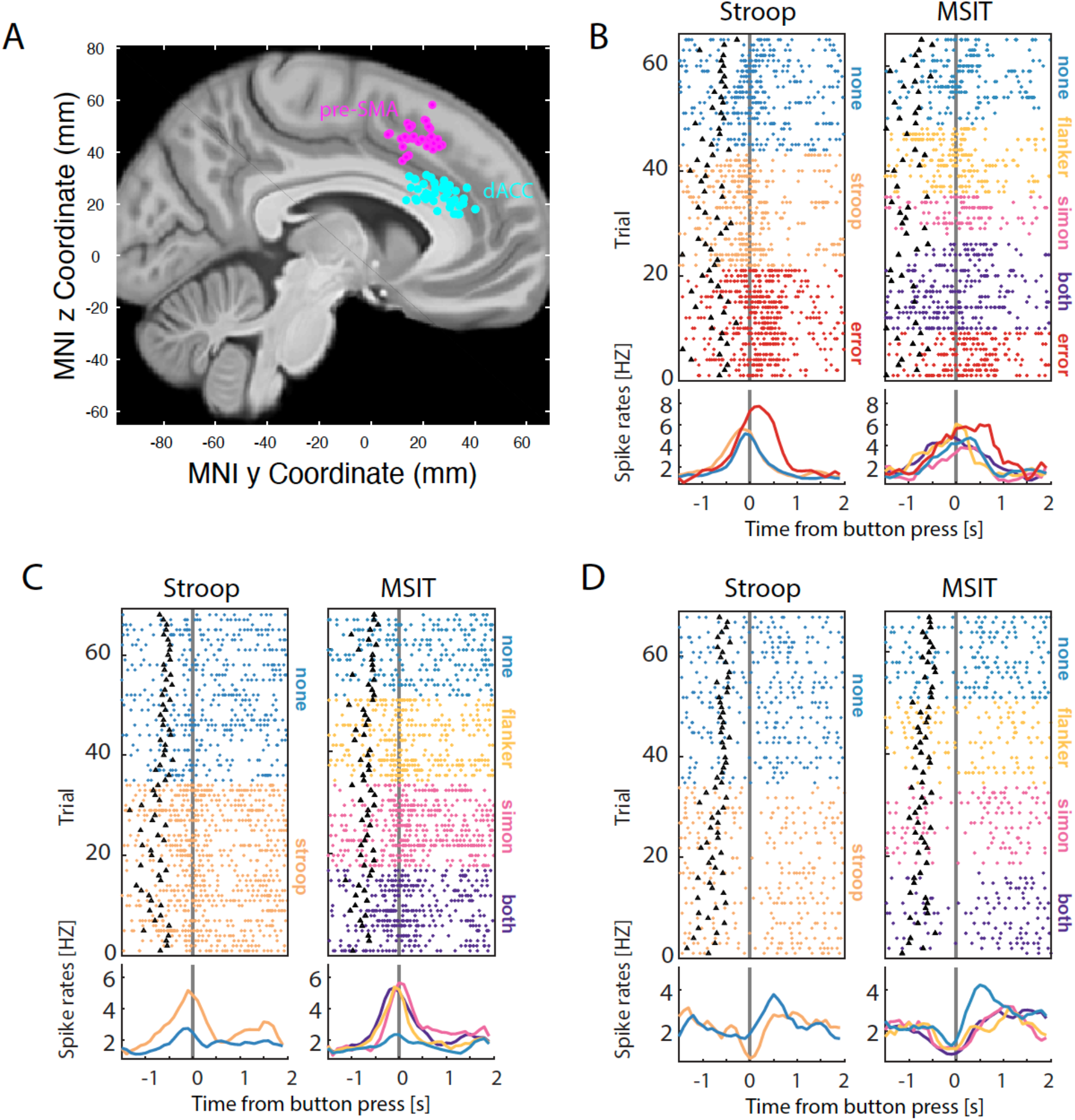
Recording locations and example neurons. (A) Recording locations shown on top of the CIT168 Atlas brain. Each dot indicates the location of a microwire bundle. (B-D) Activity of three example neurons that show similar response dynamics in both tasks. Shown is a neuron signaling action error (B), conflict by firing rate increase (C), and conflict by firing rate decrease (D). The black triangles mark stimulus onset. Trials are resorted by type for plotting purposes only.

We identified neurons selective for prior mean or prior variance in the baseline period, for conflict in the ex-ante and ex-post period, and for error, surprise, posterior, and posterior variance in the ex-post epochs (see example neurons in **Fig. 2**; schematic of analysis epochs in **Fig. 3A**; and a summary of overall cell counts in **Fig. 3B**). We pooled neurons across the dACC and the pre-SMA for most analyses because neurons responded similarly (but see section “Comparison between the dACC and the pre-SMA” for notable differences).

**Figure 3.**
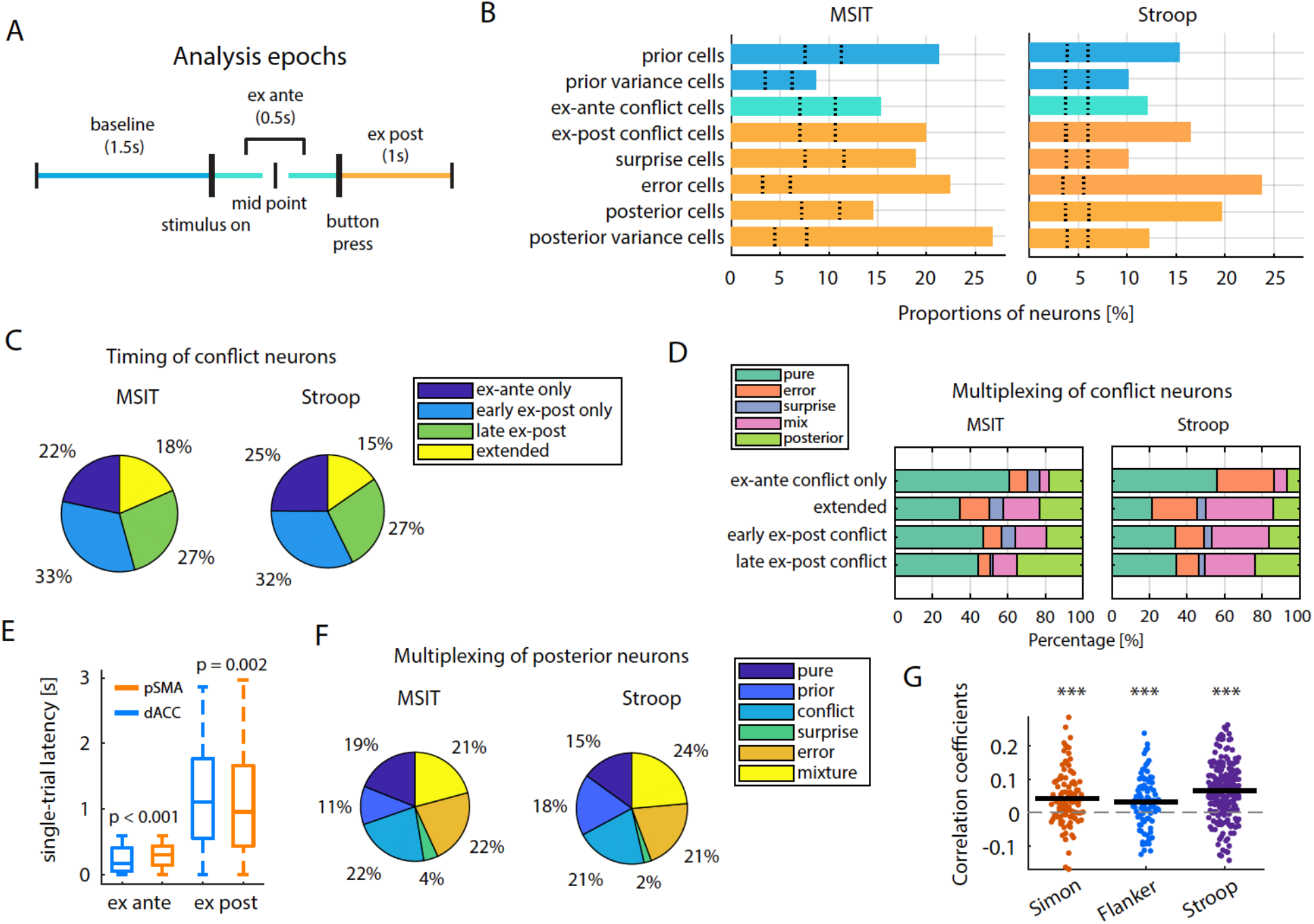
Single neuron tuning properties. (A) Illustration of epochs used for analysis. Thick vertical bars represent physical events, the slim vertical bars demarcate epochs. (B) Percentage of neurons encoding the variable indicated in the two tasks. The color code is as indicated in (A). Dotted lines represent 2.5^th^ and 97.5^th^ percentiles of the null distribution obtained from permutation. For all groups shown, p < 0.001. (C) Percentage of conflict neurons that are active in each time period. Early and late ex-post epochs denote 0-0.5s and 0.5s-1s after button press. (D) Percentage of conflict neurons that were also selective for error, surprise, posterior, or any combination of these factors (“mix”). (E) Comparison of single-trial neuronal response latency of conflict neurons in dACC and pre-SMA (correct trials only, t=0 is stimulus onset for ex-ante and button press for ex-post conflict neurons). (F) Percentage of posterior neurons that were also selective for prior, conflict, surprise, or error signaling. (G) Neuronal signature of updating conflict prior based on the posterior. Correlation is computed between the difference between prior and posterior (behavioral update) and the difference between demeaned FR_ex-post_ and FR_baseline_ (neural update) for all prior neurons.

During baseline, neurons encoded the mean or the variance of the prior distribution for conflict probability (**Fig. 3B**, blue; 18 and 13% of neurons in MSIT; 17 and 11% in Stroop). Neurons also encoded conflict ex ante (15% and 12% of neurons in MSIT and Stroop, respectively; **Fig. 3B**, green), consistent with previous reports (*34, 39*). In the ex-post epoch (**Fig. 3B**, yellow), neurons encoded conflict (20% in MSIT; 17% in Stroop), conflict surprise (an unsigned conflict prediction error generated by the experienced conflict given the conflict prior; 19% in MSIT; 10% in Stroop), errors (22% in MSIT; 19% in Stroop; see **Fig. 2A** for an example), and the mean and variance of posterior distribution of conflict probability (14/26% in MSIT; 20/12% in Stroop). The percentage of units selective for a given variable were remarkably similar between the two tasks (**Fig. 3B**).

Approximately 30% of conflict neurons were active exclusively in either the ex-ante, early (0-0.5s after button presses) or late (0.5-1.5s after button presses) ex-post epochs (**Fig. 3C;** note that ‘ex-post’ always refers to the period of 0-1s after button press unless specified as early or late), with some (~12%) active throughout the trial after stimulus onset (“extended”). This temporal distribution of conflict signals was strikingly similar between the MSIT and Stroop tasks (**Fig. 3C**). We were particularly intrigued by neurons signaling conflict ex post (15-20% of neurons in both tasks; **Fig. 2C-D** shows examples), which have not been reported before. This conflict signal, which arises too late to be useful for within-trial cognitive control, was more prominent compared to the one found in the ex-ante epoch in both tasks (15% vs 20%, χ^2^(1) = 5.08, p = 0.024 for MSIT; 12% vs 16%, χ^2^(1) = 9.19, p = 0.0024 for Stroop, chi-squared test). Motivated by our Bayesian conflict learning framework, we posit that this ex-post conflict signal serves as an “outcome” signal indicating that the trial was not only correct but also with or without conflict. As we will show further below, this information is critical for updating the conflict prior, thus dynamically adjusting the amount of cognitive control estimated for each upcoming trial.

Many conflict neurons also signaled errors, surprise, posterior, or a conjunction of these variables (**Fig. 3D**). This multiplexing of signals depended on the timing of conflict signals. The proportion of conflict neurons that multiplexed posterior information (light green bars) increased significantly towards the end of the ex-post epoch, when updating would be most complete (proportion in the late ex-post epoch vs. that in all other epochs; χ^2^(1) = 6.14, p = 0.01 for MSIT; χ^2^(1) = 6.22, p = 0.01 for Stroop, chi-squared test). Consistent with this idea, the group of neurons signaling conflict exclusively in the ex-ante epoch showed the least multiplexing, indicating a primary role in monitoring conflict during action production (proportion of “pure” conflict neurons active only during the ex-ante epoch vs. those that are active in other epochs; χ^2^(1) = 5.31, p = 0.02 for MSIT; χ^2^(1) = 8.78, p = 0.003 for Stroop, chi-squared test).

We next investigated when information about the cognitive variables identified above was available at the population level. We assessed the temporal stability of a decoder trained using data from one epoch and tested on data from a different epoch. Error decoders trained on early ex-post data generalized poorly to later epochs, but the ones trained on late ex-post data generalized well into earlier epochs (**Fig. S5A-B**), suggesting that new information emerged late in time, possibly related to post-error adjustments (*34*). We also found that the conflict coding patterns during the ex-ante epoch did not generalize well to the ex-post epoch, and vice versa (**Fig. S5C-E**), confirming that the MFC encoded these two types of conflict information with separate populations of conflict neurons. The magnitude of conflict prior (or posterior) could also be decoded from population activity at the single-trial level. Decoding conflict prior from activity in the baseline epoch generalized well to decoding conflict posterior in the ex-post epoch, demonstrating temporally stable population coding of conflict across these epochs (**Fig. S6A-F**). Representation of previous conflict was relatively weak (**Fig. S7A-C**; < 60% decoding accuracy), which is consistent with our behavioral observation that previous conflict alone was a poor predictor of RT compared to the integrated conflict prior (**Table S2**, right).

Posterior neurons demonstrated the greatest degree of multiplexing (**Fig. 3F**). Only ~18% of posterior neurons signaled posterior exclusively, with the remainder also signaling prior, conflict, surprise, or mixtures of these. This multiplexing may reflect the process of computing the posterior from the prior, which incorporates these multiple variables. Supporting this hypothesis, conflict prior-encoding neurons changed their ex-post firing rates by an amount that was commensurate with the numerical change from prior to posterior (estimated by the behavioral model) on a trial-by-trial basis (**Fig. 3G**, p < 0.001, t test against zero. Mean correlations in Simon, Flanker and Stroop are 0.042, 0.032, 0.065, respectively), reflecting a neural “updating” process.

The properties of conflict prior neurons differed from those of the other recorded neurons. First, the trial-by-trial baseline spike rates of prior neurons (a time series) had higher self-similarity than all other types of neurons studied (**Fig. S3C-D**; self-similarity quantified by α values from the Detrended Fluctuation Analysis. **Fig.S3E-H** show DFA analysis of two example neurons). Second, the spikes of prior neurons had significantly shorter action potential waveforms compared to non-prior neurons (**Fig. S3I-J** right panels; more broadly, DFA alpha is negatively correlated with spike width, shown in **Fig. S3I-J** left panels). Together, these data demonstrate that error, conflict, and conflict prior (or posterior) information can be read out reliably on each trial from the MFC population with high accuracy, with dynamic coding patterns for conflict and error and static coding patterns for prior (or posterior).

### Representational geometry of conflict within a single task

We next investigated the representational geometry of conflict when different types of cognitive conflict coexist within a single task (MSIT) in the high dimensional neural state space formed by all recorded neurons. We first tested whether the four MSIT conflict conditions are separable along a one-dimensional line in the neural state space. This “coding dimension” was defined as a vector flanked by the means of the “sf” and “none” trials (**Fig. 4A**, dotted lines for an illustration; see **Methods**). Projecting left-out single trials from all four trial types (si, fl, sf, non-conflict) onto this coding dimension allowed a decoder to differentiate between all pairs of conflict conditions in the ex-ante epoch (**Fig. 4B**, left), and all but one pair (si vs. sf, p = 0.11) in the ex-post epoch (**Fig. 4B**, right). Notably, the left-out si and fl conditions can be differentiated well on this axis (decoding accuracy is 72% for ex-ante and 78% for ex-post data). Conflict conditions were separable even after equalizing for RT across conditions (**Fig. S8C**; See **Methods** for RT equalization), suggesting that the separation was independent of trial difficulty for which RT is a proxy (*55*). We next investigated whether a common coding dimension exists for both Simon and Flanker conflict by projecting the activity of single trials onto the coding dimension formed by connecting, in the neural state space, the mean of Simon (si+sf) with the mean of non-Simon (fl+none) separately for each time bin (and vice-versa for Flanker (fl+sf) vs non-Flanker (si+none)). Data for testing were held out (not used for constructing the coding dimensions). Coding dimensions for one type of conflict allowed decoding of the other type of conflict with high accuracy (**Fig. 4C**; black trace, Flanker coding dimension decoding Simon vs. non-Simon; gray trace, Simon coding dimension decoding Flanker vs. non-Flanker). Together, these data demonstrate that within a single cognitive task, the MFC population formed a conflict representation geometry that generalized across two types of conflict while at the same time also allowing maximal separation between the different types of conflict.

**Figure 4.**
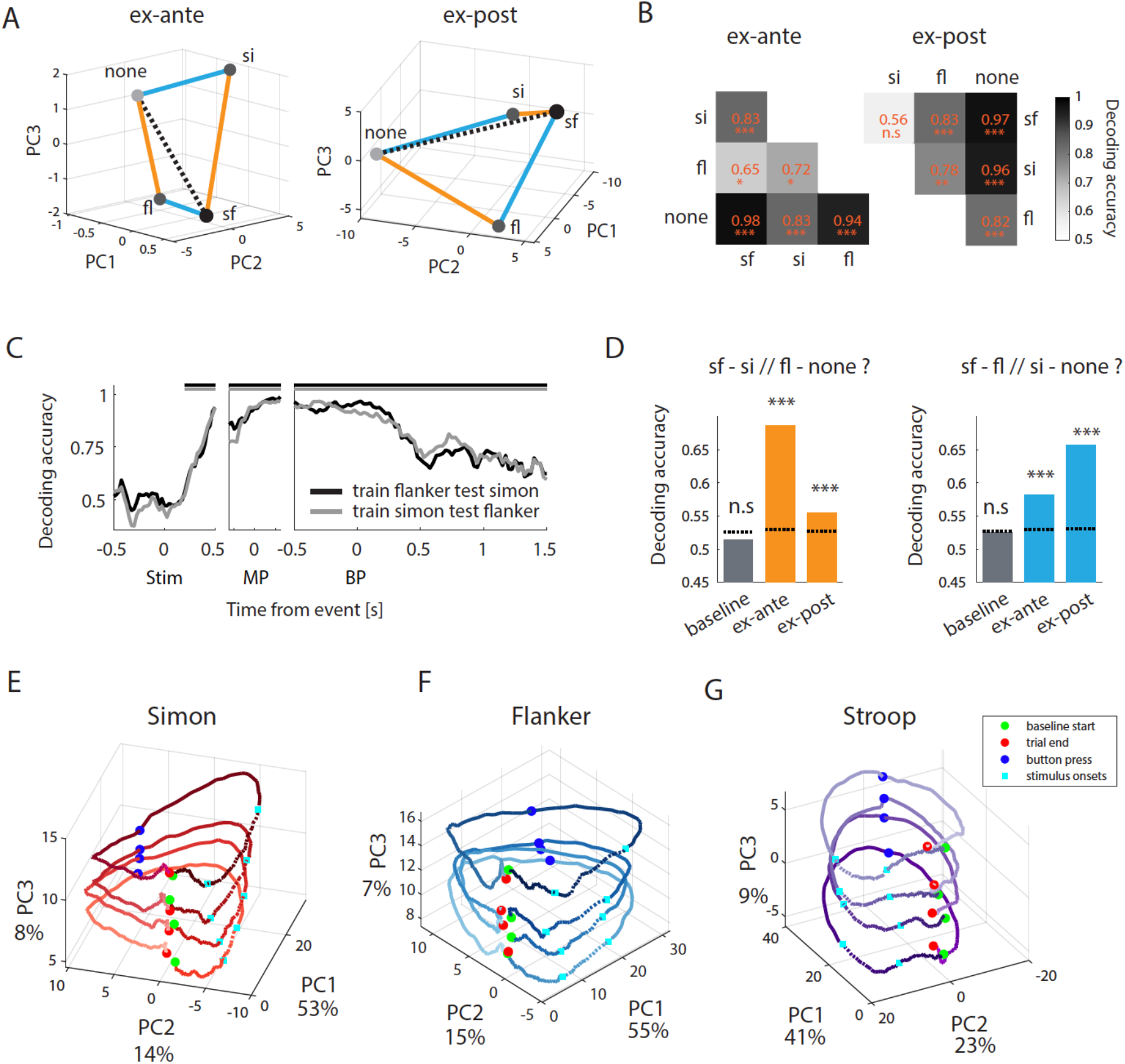
State-space representation of conflicts, prior, and posterior. (A) State-space representation of conflict (left, ex-ante; right, ex-post) in MSIT task visualized in PCA space. Dotted line is the vector used to classify pairs of conflict conditions in (B). (B) Decoding accuracy from classification of pairs of conflict conditions in MSIT. (C) Coding dimensions invariant between Simon and Flanker conflict. At each time point, a decoder is trained on Simon vs. non-Simon and tested on held out Flanker vs. non-Flanker trials (gray), and vice-versa (black). (D) Conflict representations are compositional. Decoders trained on one edge of the parallelogram were able to differentiate between conditions along the opposite parallel edge (orange and blue edges shown in A, respectively, shown on left and right). Dotted lines show 97.5^th^ percentile of the null distribution. (E-G) State-space representation of prior/posterior in MSIT and Stroop, visualized in PCA space. Green dots mark baseline start, the two cyan squares delineate the range of stimulus onsets, blue dots mark button press and red dots mark end of trial. Trials are aligned to button press. Color fades as the trial progresses.

We next tested whether the representation of conflicts was compositional. If true, relative to the mean of “none” trials, the “sf” representation should be located where the linear vector sum of Simon (“si”) and Flanker (“fl”) trials lands, forming a parallelogram with the none-sf axis as the diagonal (**Fig. 4A**). A decoder trained to differentiate between the two classes connected by one edge of this parallelogram should be able to decode the two classes connected by the opposite edge (and vice-versa). Indeed, a decoder trained to differentiate “sf” from “fl” trials, which is simply the axis connecting “sf” and “fl” (blue edge in **Fig. 4A**), was able to decode “si” from non-conflict trials projected to this axis with above chance performance, and vice versa (**Fig. 4D**, p < 0.001 for both the ex-ante and ex-post data, permutation test). The same was true for the other pair of edges (**Fig. 4D**, testing blue edges in **Fig. 4A**; p < 0.001 for both the ex-ante and ex-post data, permutation test). The parallelism was not perfect because the decoding accuracy, while well above chance, was relatively low (< 70%) compared to within-condition decoding performance (**Fig. 4C**). Neurons that encoded Simon and Flanker non-linearly (as measured by the F statistic of the interaction term between Simon and Flanker derived from an ANOVA model) contributed the most to this deviation from parallelism at the population level (**Fig. S8F**, r = 0.74, p < 0.001, for ex-post data; **Fig. S8E**, r = 0.75, p < 0.001, for ex-ante data; Spearman’s rank correlation). Thus, neurons that encoded non-linear mixtures of the two types of conflict were the cause of deviation from linear compositionality. This representation structure was disrupted on error trials: generalization performance dropped significantly in the ex-ante (**Fig. S8D**; for both edges, 68% and 58% on correct trials vs. 56% and 47% on error trials) as well as the ex-post epoch (**Fig. S8D**; for both edges, 55% and 66% on correct trials vs. 51% and 59% on error trials) on error trials. Collectively these data suggest that in the MSIT task, neural representations of conflict were structured in a compositional way that separated the four conflict conditions in a parallelogram, and this geometry was behaviorally relevant.

### Representational geometry of conflict prior/posterior

Conflict prior can be viewed as a state (an initial condition) that is present before stimulus onset and to which the population returns after completing a trial. To test this idea, we binned trials by quartiles of conflict prior (or posterior. 4 “levels” were generated) and aggregated data by these labels of conflict trial levels. PCA analysis revealed that the variability across different levels of prior/posterior (~8% of explained variance) was captured mostly by a single axis (PC3s in **Fig. 4E-G**; green dots mark trial start, red dot trial end), which was orthogonal to most of the time-dependent state changes (captured by PC1s and PC2s, ~68% of explained variance). At baseline (demarcated by green and cyan filled circles in **Fig. 4E-G**), the neural state evolved with low speed (blue points in **Fig. S8G-I**). The neural state then changed more quickly after stimulus onset (orange points in **Fig. S8G-I**; p < 0.001, paired t-test), before eventually returning to baseline near the starting position (red filled circles in **Fig. 4E-G**; yellow points in **Fig. S8G-I**). The distance between the four trajectories was thus stable across time, consistent with representation of conflict prior (or posterior) being stable states. The PCA had no access to the numerical order of conflict prior (or posterior) levels, but the projection values of neural data onto PC3 followed this order (**Fig. S8J-I**, see **Legends** for statistics of the multinomial logistic regression). Taken together, this shows that the representation of conflict prior (or posterior) information in MFC is low-dimensional, stable across time and parametric, consistent with the dynamics of line attractors.

### Domain-general performance monitoring signals at the population level

We next investigated whether the geometry of performance monitoring representations supports readout invariant across MSIT and Stroop, while simultaneously allowing maximal separation of conditions specific to MSIT. We have demonstrated above that, within MSIT, a geometry can be extracted that supports invariance across types of conflicts while keeping the four conflict conditions separated. However, it remains unclear whether this representation is task specific and only possible because of sharing a task set. Here, we studied the activity of the same neurons in two behavioral tasks to test whether representations in MFC can be independent of specific tasks sets (**Table S1** shows session information). We used demixed PCA (dPCA) (*56*) to factorize population activity into coding dimensions for performance monitoring variables (error, conflict and conflict prior (or posterior)) and task identity. The statistical significance of the extracted coding dimensions was assessed by out-of-sample decoding. The extent of “demixing” was quantified by the angle between the performance monitoring and task identity coding dimension.

The dPCA coding dimensions extracted using error and conflict contrasts (Stroop-Simon and Stroop-Flanker separately) each explained 19-21% of variance, supported task-invariant decoding and were orthogonal to the task identity dimension (**Fig. 5A** for error, **Fig. 5B** and **Fig. S9A** for conflict decoding over time, and **Fig. 5C** for conflict decoding in the ex-ante and ex-post epochs separately. p < 0.001, permutation tests; clusters with significant decoding performance demarcated by horizontal bars; For error, angle = 94.47°, p=0.53, tau=-0.032; for Stoop-Simon conflict, angle = 81.13°, p=0.19, tau=-0.048; for Stroop-Flanker conflict, angle = 78.6°, p=0.02, tau=0.086; Statistics are Kendall rank correlation to test for difference vs. 90°, “n.s.”, not significantly different from orthogonality).

**Figure 5.**
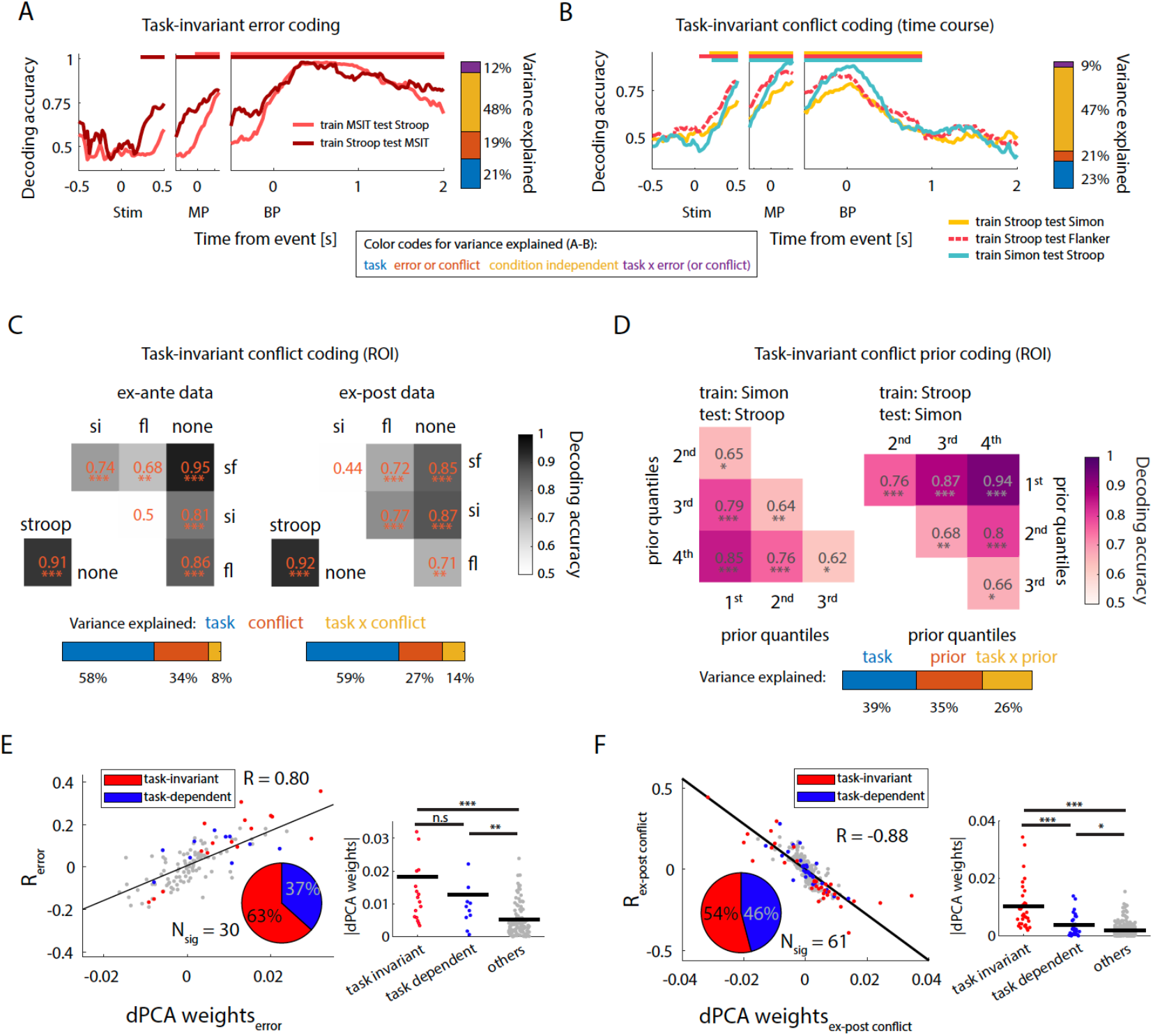
Domain-general representation of performance monitoring signals. (A) Task-invariant decoding of errors. Bar on the right shows the variance explained by the different dPCA components. (B) Task-invariant decoding of conflict. The bar on the right represents variance explained by the different dPCA components (color code see figure legend). (C) Separability of conflict conditions along the domain-general conflict axis in both the ex-ante (left) and ex-post epochs (right). The dPCA coding dimension was constructed using Stroop, SF conflict and no conflict trials and supported decoding of Stroop, Simon and Flanker conflicts (“task-invariant”) as well as separation of Simon and Flanker conditions. (D) Task-invariant decoding of conflict prior. The dPCA coding axis was constructed using Stroop and Simon conflict priors (binned by quartiles into four levels) and supported pairwise decoding of conflict prior levels in both tasks. Bar at the bottom shows variance explained of dPCA components. (E-F) Relationship between task-invariant single neuron tuning strength of error (E), conflict (F) and dPCA weights. Pie charts show the percentages of *tuned* neurons (total number is N_sig_) that had a significant main effect for performance monitoring variable *only* (“task-invariant”, red slice) and those that had a significant interaction effect with the task identity (“task-dependent”, blue slice). Scatter plots (left) shows significant correlation between task-invariant coding strength and the corresponding dPCA weights. Y-axis shows correlation of firing rate of a neuron with the given variable, after partialing out task identity (see Methods). *p < 0.05, ** p < 0.01, *** p <= 0.001, n.s., not significant (p > 0.05).

The task-invariant conflict coding dimension (extracted by removing the task difference between MSIT *sf* and Stroop conflict trials and between non-conflict trials in both tasks) was also able to differentiate 5 out of 6 pairs of conflict conditions with high accuracy (60% - 90%) within MSIT in both the ex-ante and ex-post epochs (**Fig. 5C**; statistics see figure legend, permutation tests). Task-invariant decoding did not change after RT equalization (e.g., selecting conflict and non-conflict trials that had similar RTs), suggesting again that task-invariant representations of error and conflict was not due to coincidental condition differences in difficulty (**Fig. S9B-E**), for which RT is a proxy (*55*).

Similarly, the coding dimensions for Stroop-Simon conflict prior (**Fig. 5D**; or posterior, **Fig. S9G**) and Stroop-Flanker conflict prior (**Fig. S9F**; or posterior, **Fig. S9H**) each explained 27-35% of variance, supported task-invariant decoding and were orthogonal to the task identity dimension (see Legend for statistics). Importantly, this task generalizability did not compromise the capacity of this coding dimension to separate different levels of prior (or posterior) within Stroop or MSIT (**Fig. 5D**, **Fig. S9F-H**). Together, these data demonstrate that the neural representation of performance monitoring signals in MFC is configured in such a way that it supports generalization between two different tasks while at the same time also allowing the readout of task-specific information.

### Domain-general performance monitoring signals at the single-neuron level

We sought to understand how single neuron tuning properties contributed to the population geometry that simultaneously allowed task-invariant and task-specific readouts. While some neurons encoded error, conflict, and conflict prior (or posterior) in a task-invariant manner (**Fig. 2**), others encoded these variables in only one task. We assessed the level of task specificity with a linear regression model with two main effects: the task identity and one of the performance monitoring variables (see **Methods**). 33-66% of the recorded neurons were “task invariant” as assessed by having a significant main effect of error, prior, conflict prior (or posterior) but no significant interaction with task identity (**Fig. 5E-F, Fig. S11A-E**, red part of pie charts). 37-63% of the recorded neurons were “task dependent” as assessed by a significant interaction effect (**Fig. 5E-F**, **Fig. S11A-E**, blue part of pie charts). Both the task-invariant and the task-dependent neurons contributed to the task-invariant coding dimensions as both were assigned significantly larger absolute dPCA weights compared with non-selective (“other”) neurons (**Fig. 5E-F**, **Fig. S11A-E**, dot density plots). The dPCA weight assigned to each neuron was correlated with the neuron’s task-invariant single-neuron coding strength (scatter plots in **Fig. 5E-F**, **Fig. S11A-E**; p < 0.001 for all panels, Pearson’s correlation). Neurons with higher self-similarity for baseline spike counts contributed more to the task-invariant coding dimension of conflict prior (or posterior), after controlling for single-neuron coding strength (partial correlation between absolute values of dPCA weights and DFA α values; r = 0.22 and r = 0.24, p < 0.001 for Stroop-Simon prior and posterior; r = 0.16, p = 0.003 and r = 0.16, p = 0.002 for Stroop-Flanker prior and posterior). These results reveal that a downstream neuron receiving input from a random subset of MFC neurons can robustly extract domain-general performance monitoring signals with a set of properly tuned connection weights.

### Comparison between the dACC and the pre-SMA

The proportion of neurons selective for each tested variable were similar in dACC and pre-SMA (**Fig. S3A-B).** Both areas contained similar proportions of task-invariant and task-dependent neurons. Both areas independently supported compositional conflict coding and domain-general readouts for error (**Fig. S10A-B**), conflict (**Fig. S10C-F**), and conflict prior (or posterior; **Fig. S10G-N**). However, there were three notable differences between the two areas. First, the temporal profiles of decoding performance for error, conflict and prior (or posterior) were similar between the areas but decoding accuracy in pre-SMA was consistently higher (**Fig. S4A-E**). Second, the task-invariant error response appeared earlier in pre-SMA than in dACC (Δ_latency_ = 0.55s for Stroop and 0.5s for MSIT), consistent with our previous report in the Stroop task (*34*) (replicating this difference for MSIT is novel). Third, ex-ante conflict information was first available in dACC, followed by pre-SMA (**Fig. 3E**; median difference = 138ms; p < 0.001, Wilcoxon rank sum test; single-trial spike train latency). Third, by contrast, ex-post conflict information was available first in pre-SMA, followed by dACC (**Fig. 3E**; median difference = 161ms; p = 0.002, Wilcoxon rank sum test). This pattern is consistent with a leading role of pre-SMA in post-action performance monitoring (*34*), and that of dACC in conflict monitoring during action selection.

## Discussion

We show that some neurons encoded error, conflict and conflict prior (or posterior) in a task-invariant way, some encoded these variables exclusively in one task only, and some multiplexed domain/task information to varying degrees, intermixed at similar anatomical locations within the MFC. Thus, at the single neuron level, neurons multiplex task identity and performance monitoring variables nonlinearly (i.e., “mixed selectivity” (*47, 48*)), making simple interpretation of domain generality impossible. Population activity, however, could be factorized into a task identity dimension and a task-invariant dimension that were orthogonal to each other. The geometry of MFC population activity allows linear decoders to read out performance monitoring variables with the same high accuracy (>80%) across tasks, and simultaneously to discern different types of conflicts or different levels of conflict prior (or posterior) within a task. Importantly, it was the *same* group of neurons that gave rise to this geometry. This contrasts with neuroimaging studies that either report domain-specific and domain-general conflict signals encoded by distinct groups of voxels (*57, 58*), or an absence of domain-general conflict signals (*59*) within the MFC.

Several influential studies have proposed that the lateral PFC is topographically organized to subserve cognitive control, with more abstract processing engaging the anterior regions (*1–3, 60, 61*). Unlike in the LFPC, we find that domain-general and domain-specific neurons are intermixed within the MFC. The representational geometry we report here is well suited to provide performance monitoring signals to these subregions: downstream neurons within these LPFC regions can select performance monitoring information at different degrees of abstraction by adjusting connection weights, similar to input selection mechanisms described in the PFC (*62*). Curiously, we did not observe a prominent difference between the pre-SMA and the dACC in the degree of domain generality of performance monitoring.

### Domain-general error signals

A key component of performance monitoring and metacognitive judgement is the ability to detect action errors without relying on external feedback (*63, 64*). Here, we show that a subset of neurons not only signal error in the Stroop task (as previously reported in (*34*)) but also in the MSIT (a novel finding). The error signal is thus domain-general, that is, abstracted away from the sensory and motor details as well as the types of response conflicts encountered across the two tasks (**Fig. 2B**). Remarkably, at the population level, these domain-general error neurons enabled trial-by-trial readout of self-monitored errors with > 90% accuracy equally across tasks (**Fig. 5A, E**). Such abstract action error signals, which indicate a mismatch between abstract goals and action outcomes, may provide a basis for the processes that monitor the difference between internal thoughts and reality, which are thought to be compromised in psychiatric diseases such as schizophrenia (*65*). Given the fMRI-BOLD finding that the MFC is a domain-general substrate for metacognition (*66*), an interesting open question is whether the same neural mechanisms we describe here support metacognitive judgement across different domains, such as perceptual or memory confidence.

### Domain-specific performance monitoring

The causes of conflict and errors in the two tasks differs: distraction by the prepotent tendency to read in the Stroop task, and distraction by location of target number (Simon) or by numbers flanking the target (Flanker) in the MSIT. These performance perturbations call for specific compensatory mechanisms, such as suppressing attention to task-irrelevant stimulus dimensions. Consistent with this requirement, a subset of neurons signaled errors, conflict and conflict prior (or posterior) exclusively in one task. At the population level, these neurons contributed to successful decoding of the different types of conflicts within MSIT and gave rise to a task identity dimension that supported robust decoding of which task the performance disturbance occurred in (>90% accuracy), providing the domain-specific information about the sources of performance disturbances for cognitive control. These results are broadly consistent with the reported role of MFC neurons in credit assignment (*17*). The existence of task-specific neurons also suggests that the performance monitoring circuitry can be rapidly and flexibly reconfigured in different tasks to subserve different task sets (*67, 68*), which is consistent with the rapid reconfiguration of functional connectivity among cognitive control networks to enable novel task performance (*69*).

### Compositionality of conflict representation

Further insight into how representations can be both general and specific is offered by examining the activity during “sf” conflict trials (both Simon and Flanker conflict) in the MSIT task. The problem of whether conflict representations are compositional can be formulated as a problem of cross-condition generalization: if Simon and Flanker conflict are linearly additive, decoders trained to identify the presence of only Simon or Flanker conflict should generalize to trials in which both types of conflict are present. We found that this was the case, with the neural state approximately equal to the linear vector sum of the two neural states when the two types of conflict are present individually. Conflict representations are thus additive to a large extent (with the extent of deviation predicted by the degree of nonlinear mixing present). The (approximate) factorization of conflict representation is important for both domain-specific and domain-general adaptation: when different types of conflict occur simultaneously and the representation can be factorized, downstream processes responsible for resolving each type of conflict can all be initiated. On the other hand, domain-general processes can also read out the representation as a sum and initiate domain-general adaptations.

### Estimating control demand enabled by ex-post conflict neurons

Our Bayesian updating model predicts that conflict should be signaled twice: once during response competition (*ex ante*), and again after the action has been committed (*ex post*). While the former is a major prediction of conflict monitoring theory (*13, 70*), the latter is a new prediction of our model. Importantly, *separate* groups of neurons gave rise to these two types of conflict signals (**Fig. 3C**) and the population activity patterns representing the two differ substantially to prevent generalization (**Fig. S5C-E**), suggesting that the ex-post conflict signal is not a continuation of the ex-ante conflict signal but instead a re-representation of experienced response conflict by different neurons. We note that the ex-post conflict signal we documented appears before onset of feedback and thus reflects metacognitive judgements. Our work is the first to report ex-post conflict neurons in the dACC and pre-SMA in both tasks and ex-ante conflict neurons in the pre-SMA. Also, our work replicates the finding of ex-ante conflict neurons in the dACC (*39*). The ex-ante conflict signal may be useful for recruiting online cognitive control (*70*), but provides an temporally unstable readout of the experienced conflict: it intensifies before ultimately becoming subdued as one response option wins over. The ex-post conflict signal, by contrast, abstracts over this temporal dynamics and integrates it into a stable “conflict outcome” (*13, 51*). This view of the ex-post conflict signal as an outcome signal is also consistent with the widely reported role of MFC in representing outcomes (*4, 37*). This information is imperative for updating slowly varying representations of estimated conflict probability.

Both conflict signals arise independent of external feedback, thus qualifying as correlates of metacognitive self-monitoring (*63, 64*). Interestingly, there is significant overlap between error neurons and ex-post conflict neurons. Confirming this, we found common coding dimensions in each task that supports the decoding of both error and conflict, though the decoding accuracy is significantly lower for conflict than for error (**Fig. S8A-B**). The ex-post conflict and error signals may be generated by a common process that compares a corollary discharge signal with the sensory prediction of action outcomes generated by a cognitive “forward model” (*71–73*) and may underlie the sense of agency (*74*). Future work is needed to test this new hypothesis.

The neurons that reported conflict prior changed their firing rates trial-by-trial as predicted from our behavioral model, and this “updating” in firing rates occurred after an action was performed. The distinct functional properties of conflict prior neurons, i.e. significantly narrower extracellular spike waveforms and higher self-similarity (autocorrelation) of trial-by-trial baseline spike counts assessed by DFA (**Fig. S3C-J**), suggest that they are likely putative interneurons with strong recurrent connectivity, consistent with a prior report where dACC neurons that encode past outcomes have narrower extracellular waveforms (*75*). At the population level, these single-neuron properties contribute to the formation of a task-invariant line attractor dynamics that stably maintains the conflict prior. Computational modelling demonstrates the importance of inhibitory interneurons in maintaining information in working memory, which occurs on the scale of several seconds(*76*). Similar circuit-level mechanisms could provide a basis for retaining the history of performance monitoring and reward, which is on the scale of several minutes. These prior neurons might thus provide a neural substrate for proactive control and learning of value of control, both of which requires stable maintenance of learned information (*13, 77*). Future research should address the trade-off between flexible updating and stable maintenance of performance monitoring information. Together, the neuronal responses we found fit remarkably well to the parameters of a Bayesian model that used trial-wise updating to estimate upcoming conflict – a critical ingredient in the control of flexible behavior in changing environments.

Our findings will also be relevant to a mechanistic understanding of impaired control, such as may happen in children, under fatigue or substance influence, or in psychiatric disorders such as schizophrenia. Given our clear evidence both for domain-general and domain-specific representations, an intriguing future direction would be to ask whether impaired control may affect one of these differentially and could be compensated differentially.

## Supplementary Figures

**Figure S1.**
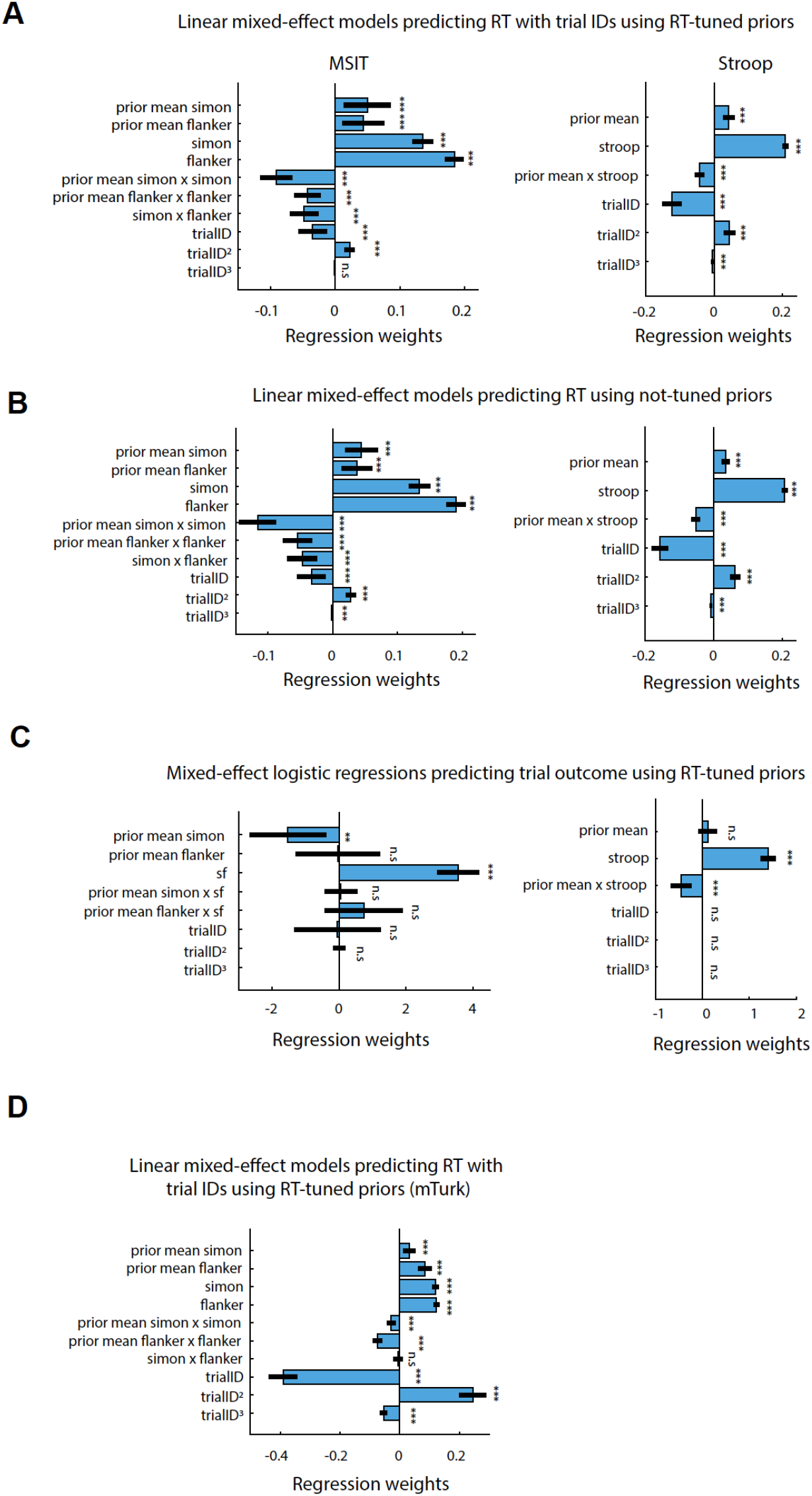
Behavioral models. Related to Figure 1. Statistical significance of regressors is determined by comparing the full model and a reduced model with a particular regressor removed, using a likelihood ratio test. (A) Linear mixed-effect model for RT that incorporates trial ID regressors for MSIT (left) and Stroop (right). Conflict priors used are from the Bayesian online learning models with RT tuning. We added the first, second and third-order trial ID regressors to model putative practice effects. The main effects of conflict priors, conflict, and their interaction are all significant even in the presence of trial ID regressors, suggesting these regressors capture behavioral effect that do not depend on trial ID. (B) Same as (A), but for the Bayesian online learning models without RT tuning. Thus, in this instance, conflict prior is estimated based on conflict sequence alone. The main effects of conflict priors (not tuned by RT), conflict, and their interaction are all significant in the presence of trial ID regressors. Therefore, RT tuning improves conflict prior (see (A) and Table S2, S3), but this is not required. (C) Mixed-effect logistic regression for predicting trial outcome (error or correct) for MSIT (left) and Stroop (right). Conflict priors used are from the Bayesian online learning models with RT tuning. For MSIT, we consider only “sf” trials for conflict trials, on which most of errors occur, and non-conflict trials. Conflict prior reduces error likelihood in both MSIT (significant main effect, p = 0.009) and Stroop (significant interaction term), p < 0.001). (D) Behavioral analysis for data collected from independent control group in the MSIT task (online experiment, see methods). Same model as used in (A) is used.

**Figure S2.**
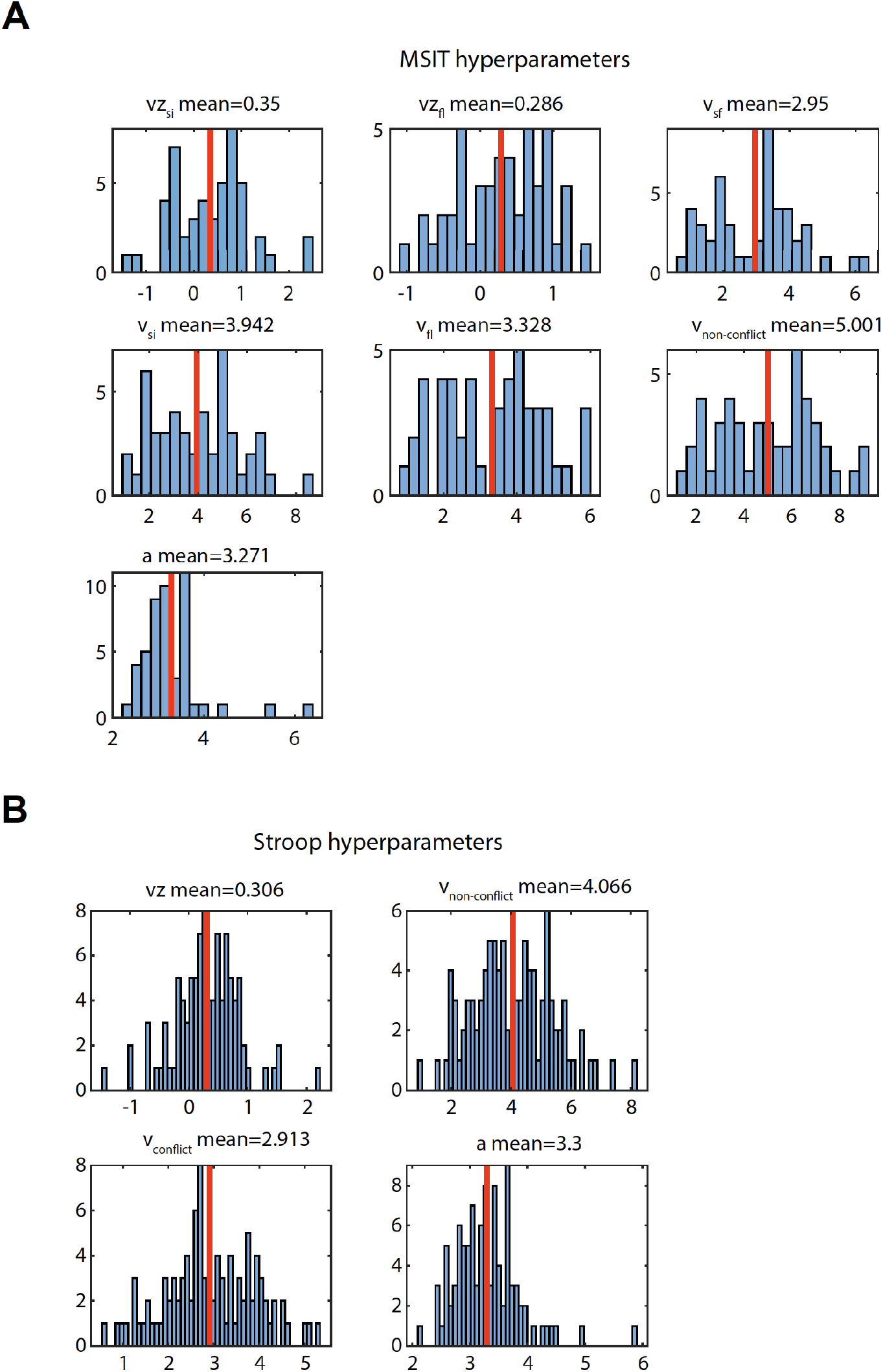
DDM hyperparameters used in Bayesian conflict learning models. Related to Figure 1. (A) Hyperparameters used in the Bayesian conflict learning model for MSIT. *vz_si_* and *vz_fl_* are coefficients scaling Simon and Flanker prior. *v_sf_, v_si_*, *v_non-conflict_* are base drift rates in both Simon and Flanker present (“sf”), Simon-only (“si”), Flanker-only (“fl”), non-conflict trials. *a* is the boundary separation. The effective drift rate was the sum of the base drift rate and the scaled conflict prior. The base drift rates were significantly different from each other (p < 0.001, ANOVA). Post-hoc pairwise testing from a multiple comparison test determined that *v_si_* did not differ significantly from *v_fl_*; *v_sf_* were significantly larger than either *v_si_* or *v_fl_*; both *v_si_* and *v_fl_* were significantly larger than either *v_non–conflict_*. (B) Hyperparameters used in the Bayesian conflict learning model for Stroop. *vz* is the coefficient scaling Stroop conflict prior. *v_non-conflict_, v_non-conflict_* are base drift rates in conflict and non-conflict trials. *a* is the boundary separation. *v_non-conflict_* are significantly larger than *v_conflict_* across sessions (p < 0.001, t test). Hyperparameters are used in the DDM likelihood function for tuning the conflict prior estimation process using an expectation-maximization algorithm.

**Figure S3.**
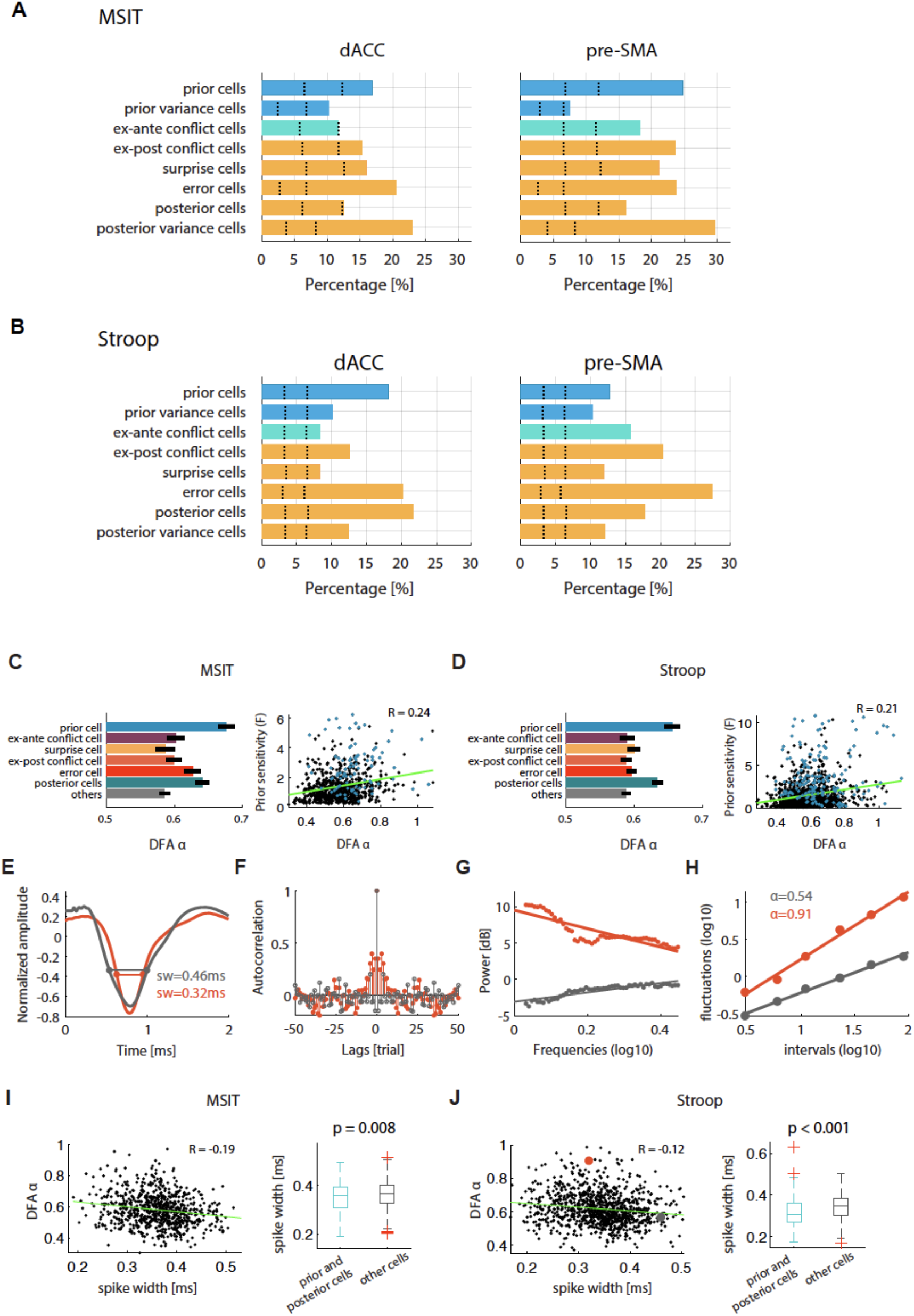
Neuronal selection by areas and long-range temporal correlation of prior/posterior neurons. (A) Percentages of significant neurons in both dACC (left) and pre-SMA (right) in MSIT. (B) Percentages of significant neurons in both dACC (left) and pre-SMA (right) in Stroop. Dotted lines represent 2.5^th^ and 97.5^th^ percentiles of the null distribution obtained from permutation. For all groups shown, p < 0.001. Patterns of neuronal selection are similar between dACC and pre-SMA. (C) Mean baseline DFA a value for different groups of neurons (left), and correlation between baseline DFA a value and the coding strength of prior, which is assessed by the F statistic computed from regressing prior against baseline spike counts (right) for MSIT. Separate data were used to compute a value and prior coding strength to avoid selection bias. Prior neurons have significant larger baseline DFA a value than any other groups (p < 0.001, ANOVA). The coding strength of prior was correlated strongly with the baseline a value (r = 0.24, p < 0.001). (D) same as in (C) but computed for the Stroop task. (E-H) Two example neurons (one orange one gray), showing waveforms (E), autocorrelation (F), power spectrum (G), and fluctuation (H) s as a function of time intervals used to compute DFA a value (slope). The neuron with narrower spike width has higher DFA a value (r = −0.19, p < 0.001). (I) DFA a value is negatively correlated with spike width for MFC neurons (left) in MSIT. Prior and posterior neurons as a group have significantly narrower spikes than other neurons (right). (J) same as in (I) but for the Stroop task.

**Figure S4.**
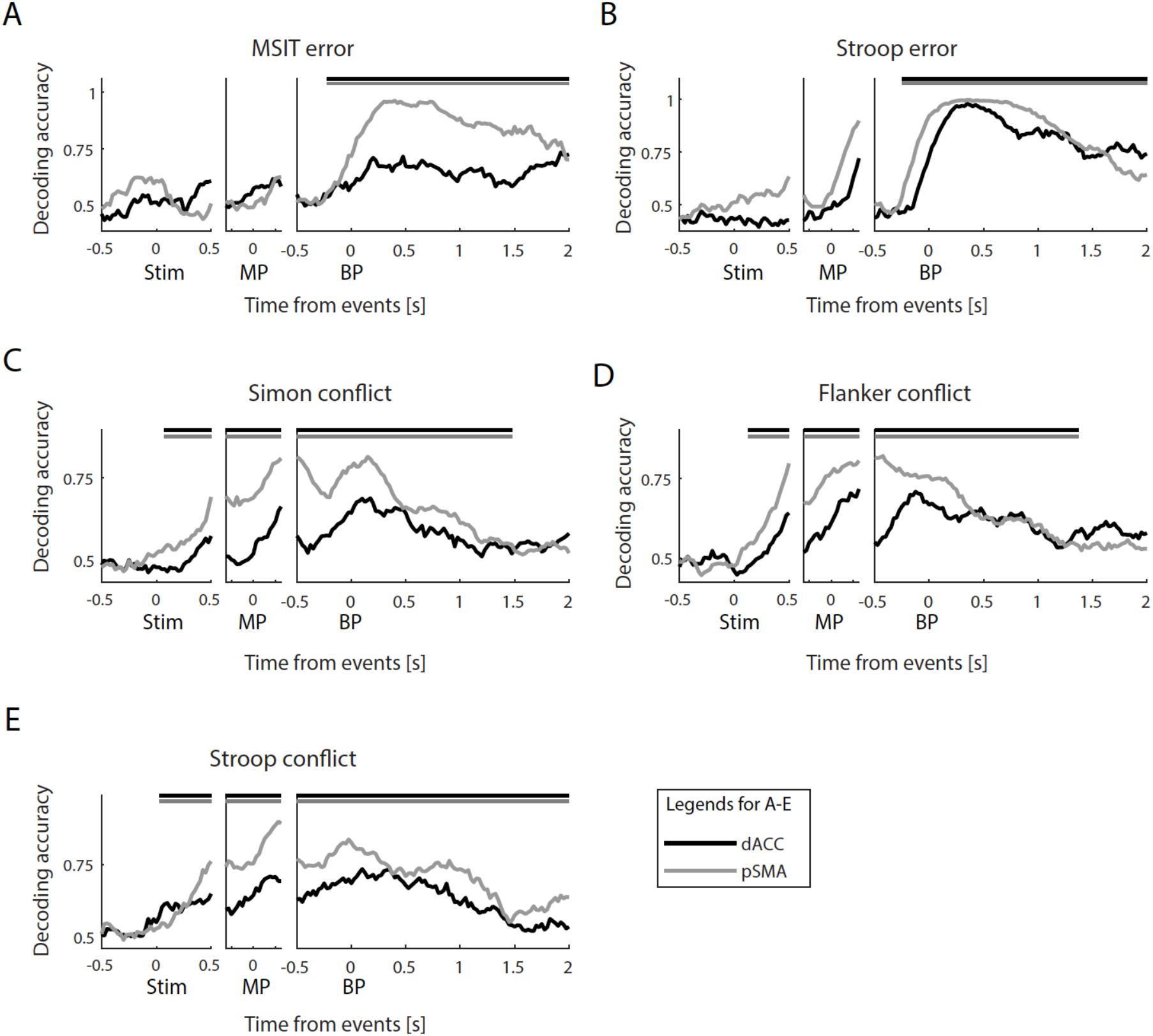
Population decoding of error and conflict by areas. (A-E) Population decoding accuracy for MSIT error (A), Stroop error (B), Simon conflict (C), Flanker conflict (D), Stroop conflict (E). For (A-E), black traces are from dACC data and grey traces are from pre-SMA data. Horizontal bars at the top demarcate significant cluster as determined by the cluster-based permutation test (p < 0.05). Overall dynamics are similar between dACC and pre-SMA, though the decoding accuracy on average is lower in the dACC.

**Figure S5.**
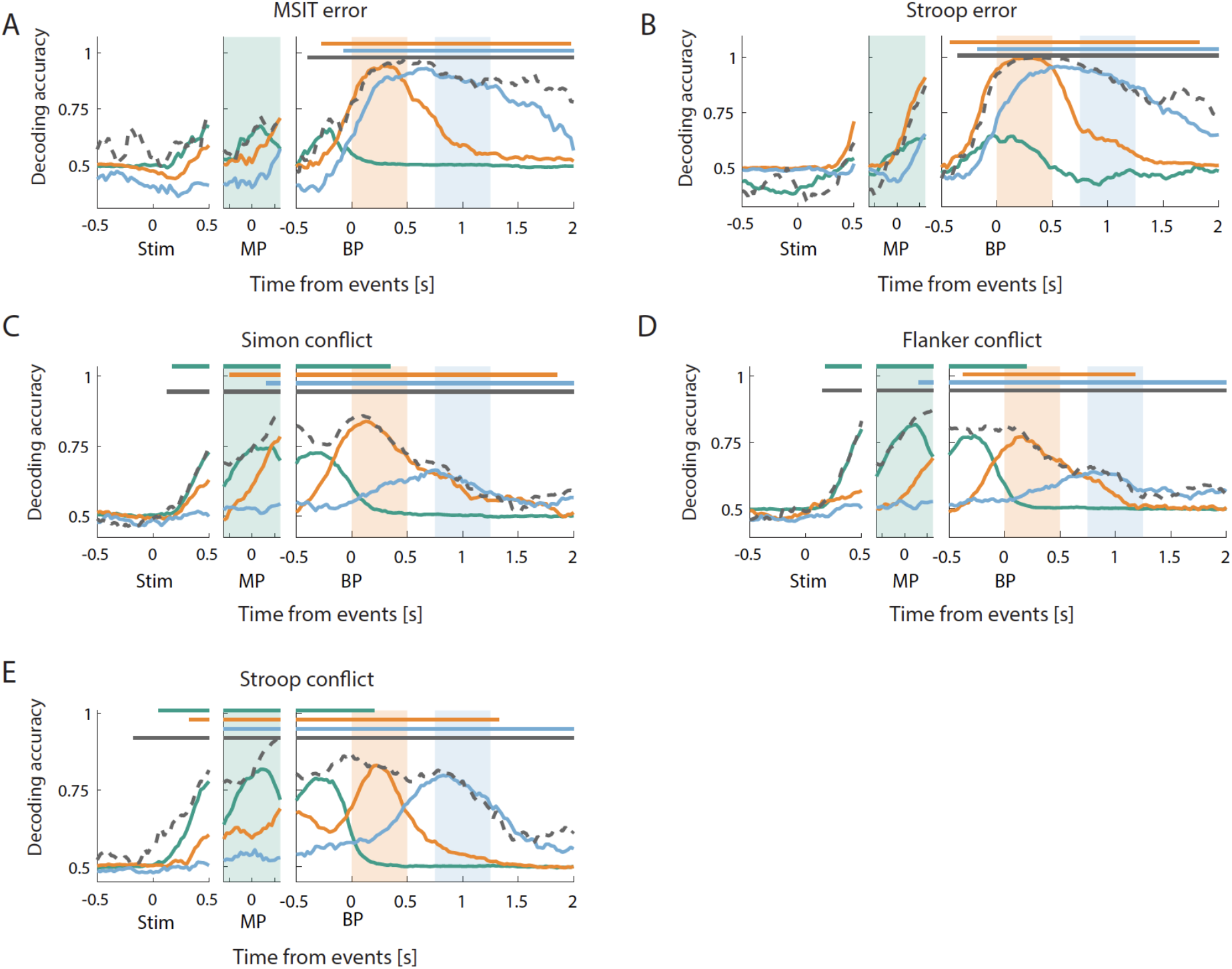
Temporal dynamics and cross-temporal generalization of error and conflict. (A-E) Decoding accuracy as a function of time for MSIT error (A), Stroop error (B), Simon conflict (C), Flanker conflict (D) and Stroop conflict (E). The three panels show data aligned to stimulus onset (“Stim”), midpoint between 100ms after stimulus onset and button presses (“MP”), button press onset (“BP”). Dotted gray trace represents within-time decoding accuracy, i.e., the data from the same epoch were used to train and test a decoder. Green, blue and orange traces represent decoding accuracy of decoders trained with data from the epochs demarcated by shading with the same colors, which is a test of the cross-temporal generalization of these decoders. Horizontal bars demarcate the extent of significant clusters in time as determined by the cluster-based permutation test (p < 0.5).

**Figure S6.**
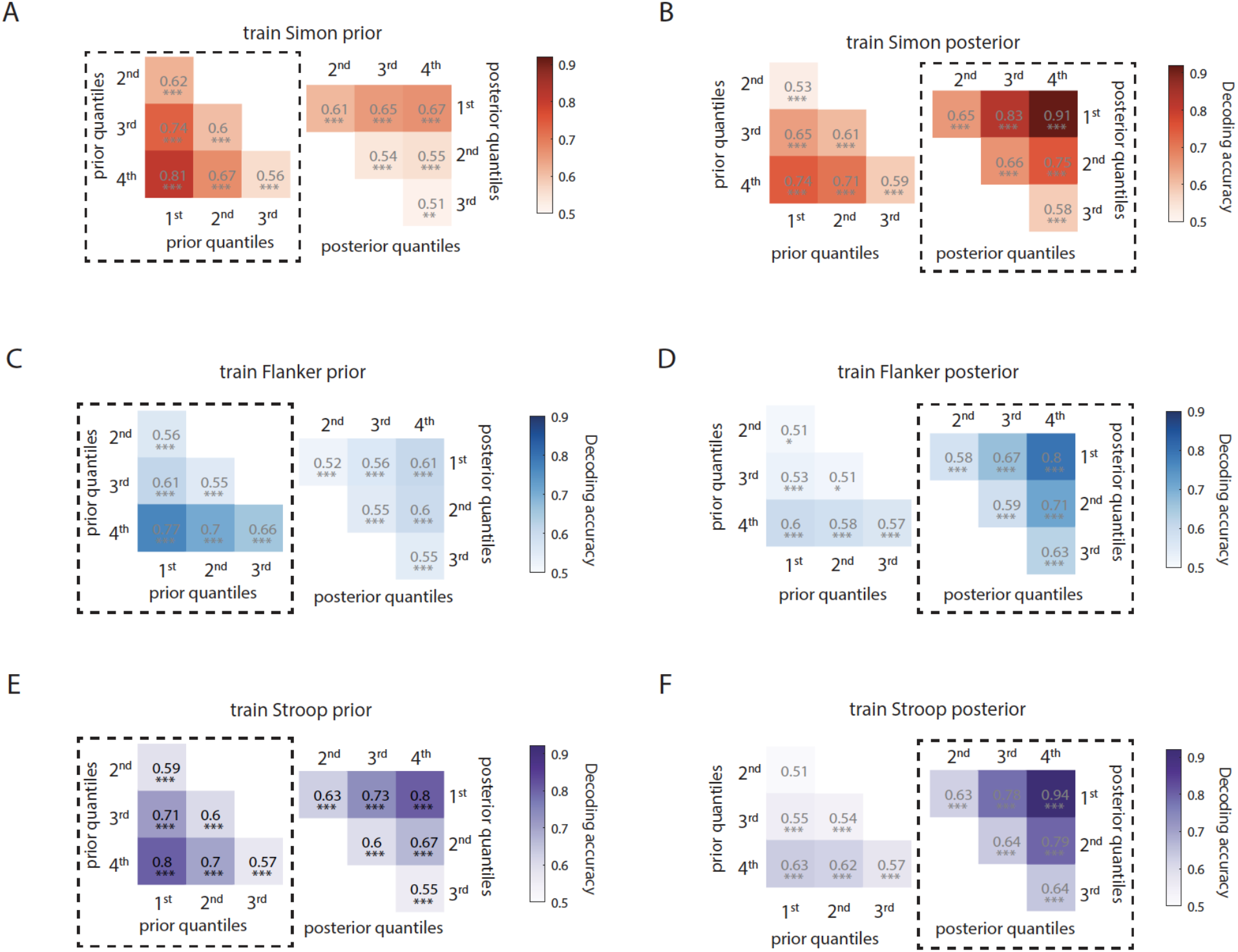
Temporal dynamics and cross-temporal generalization of conflict prior (or posterior). (A-F) Decoding accuracy for classifying between pairs of Simon (A-B) Flanker (C-D) and Stroop (E-F) prior or posterior levels. Levels are quartile bins of the conflict prior. Color bars show decoding accuracy. Dotted frames mark the within-time decoding results, plots without dotted frames showed results from temporal generalization. Decoders that classify prior levels are trained using baseline spike counts (1.5s before stimulus onset), whereas decoders that classify posterior levels are trained using ex-post spike counts (2s after button press). To test temporal generalization of these decoders, prior-trained decoders are tested with posterior data and labels, and vice versa. Dashed boxes represent within-time decoding.

**Figure S7.**
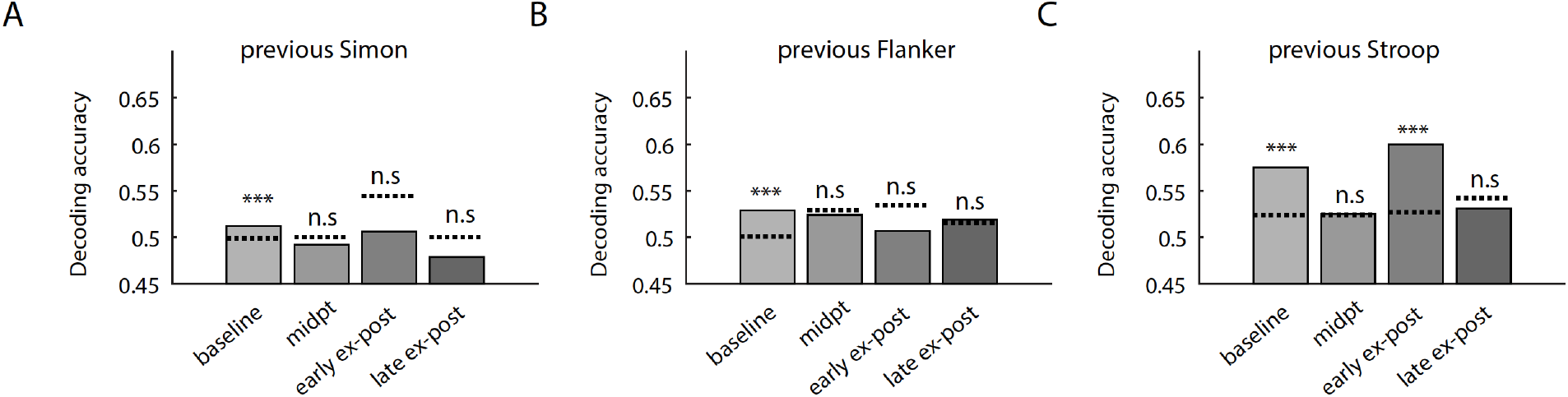
Encoding of previous conflict. (A-C) Population decoding of Simon (A), Flanker (B), Stroop (C) on the immediately preceding trial in different epochs. Dotted lines show 97.5% percentile from the null distribution (permutation). During baseline, there is significant coding of past trial conflict as expected from the persistence of ex-post conflict signals. Coding of the past trial conflict was non-significant during all other epochs except for past trial Stroop conflict in the early ex-post epochs. *p < 0.05, ** p < 0.01, *** p < 0.001, n.s., not significant (p > 0.05 or not significant determined using FDR).

**Figure S8.**
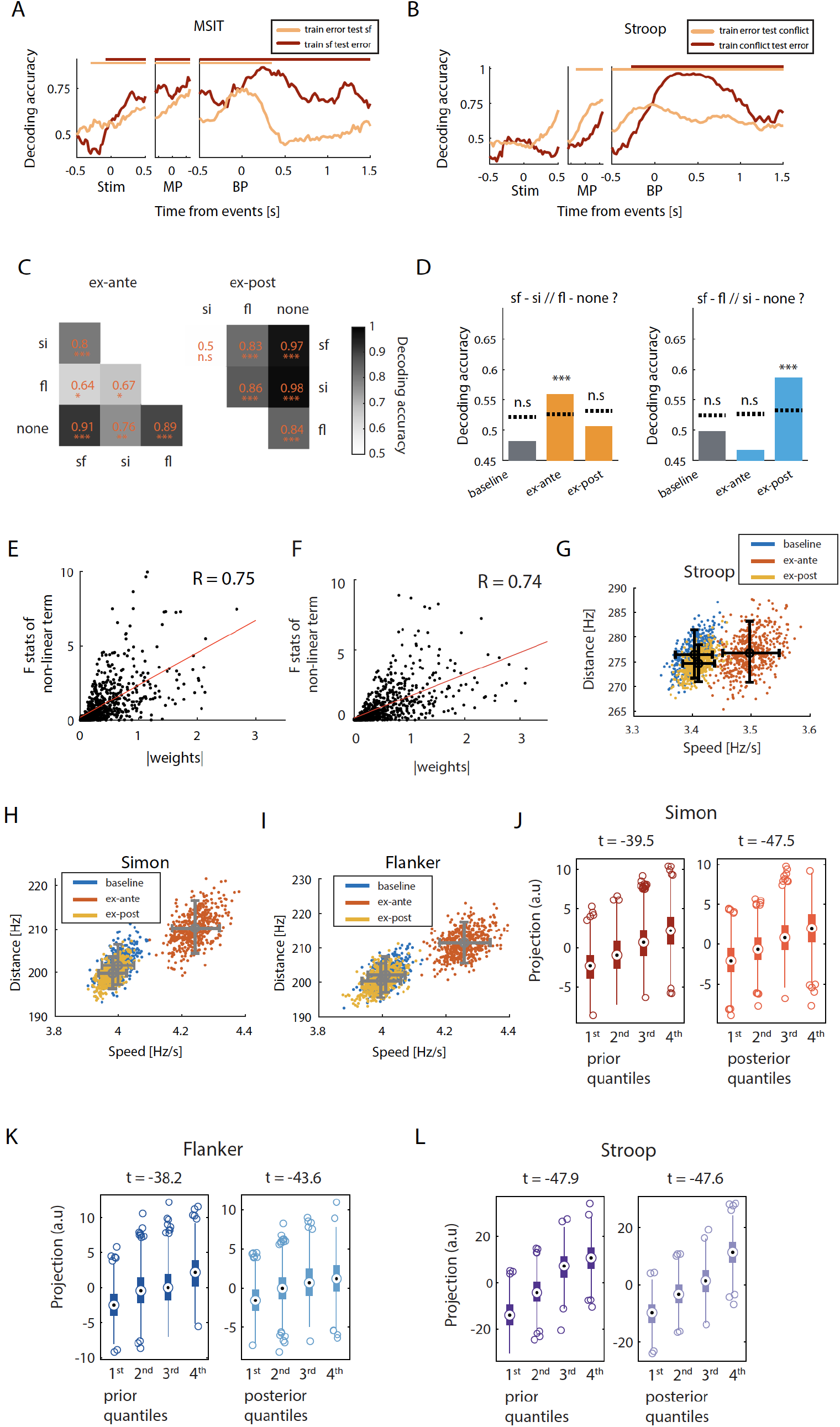
Common coding dimensions for error and conflict and MSIT state space analyses. (A) A common population coding dimension for error and conflict in MSIT. This coding dimension is extracted using dPCA, using an error contrast (error “sf” trials vs. “sf” trials) and a conflict contrast (“sf” trials vs. no conflict). “sf” trials are split into two non-overlapping groups for this. Plot show the decoding accuracy of both sf (apricot) vs no-conflict trials, and error “sf” vs. “sf” trials (out-of-sample). Horizontal bars at the top demarcate significant clusters, as determined by the cluster-based permutation test (p < 0.05). (B) Same as in (A) but for Stroop data. (C) Decoding accuracy of pairwise classification of conflict conditions after RT was equalized across conditions. Trials were selected such that RTs on si, fl, sf and non-conflict trials were equalized (p > 0.1, t test). Training data and left-out testing data from the four conflict conditions are projected to the population vector flanked by averages of non-conflict trials and sf trials, shown as the dotted line in Figure 4A. Color code represents decoding accuracy. This coding dimension separates the four conflict conditions well. (D) Testing compositionality of conflict representation with condition generalization of decoding on *error* trials. Decoding accuracy were reduced on error trials compared to on correct trials (compare with Figure 4D). Dotted lines show 97.5^th^ percentile of the null distribution from permutation. (E) Single neurons with nonlinear mixed-selectivity of Simon and Flanker conflict contribute to deviation of conflict representation from perfect linearity. Data used here are from the ex-ante epoch in (E) and ex-post epoch in (F). Nonlinear mixed-selectivity is measured by the F statistic of the interaction term between Simon and Flanker conflict in an ANOVA model with spike counts as the dependent variable. Each neuron’s contribution to the deviation from linear additivity in the high dimensional neural space is quantified by the weight of the difference vector between “sf” and “si + fl”. Scatter plot shows the relation between these two measures. Red line shows the linear fit. (G-I) Distance between trajectories and average speed computed from trials binned by quartiles of Stroop (G), Simon (H), Flanker (I) conflict prior in the baseline (blue) and the ex-ante (orange) epoch, and trials grouped by Simon conflict posterior in the ex-post epoch (yellow). Trajectories are visualized in Figure 5E-G. The state space speed stays low during baseline, increases significantly during the ex-ante epoch and decreases back to a value similar to that during the baseline. Distance between trajectories is stable across time. (J-L) Projection values on the coding dimension of Simon (J, p < 0.001 for both prior/posterior, prior t(11996) = −39.5, posterior t(11996)=-47.5), Flanker (K, p < 0.001 for both prior/posterior, prior t(11996) = −38.2, posterior t(11996)=-43.6) or Stroop (L, p < 0.001 for both prior/posterior, prior t(11996) = −47.9, posterior t(11996)=-47.6) prior or posterior (PC3 in Figure 5E-G). The order of projection values is on average consistent with the order of the prior/posterior levels, even though PCA does not have access to the order information.

**Figure S9.**
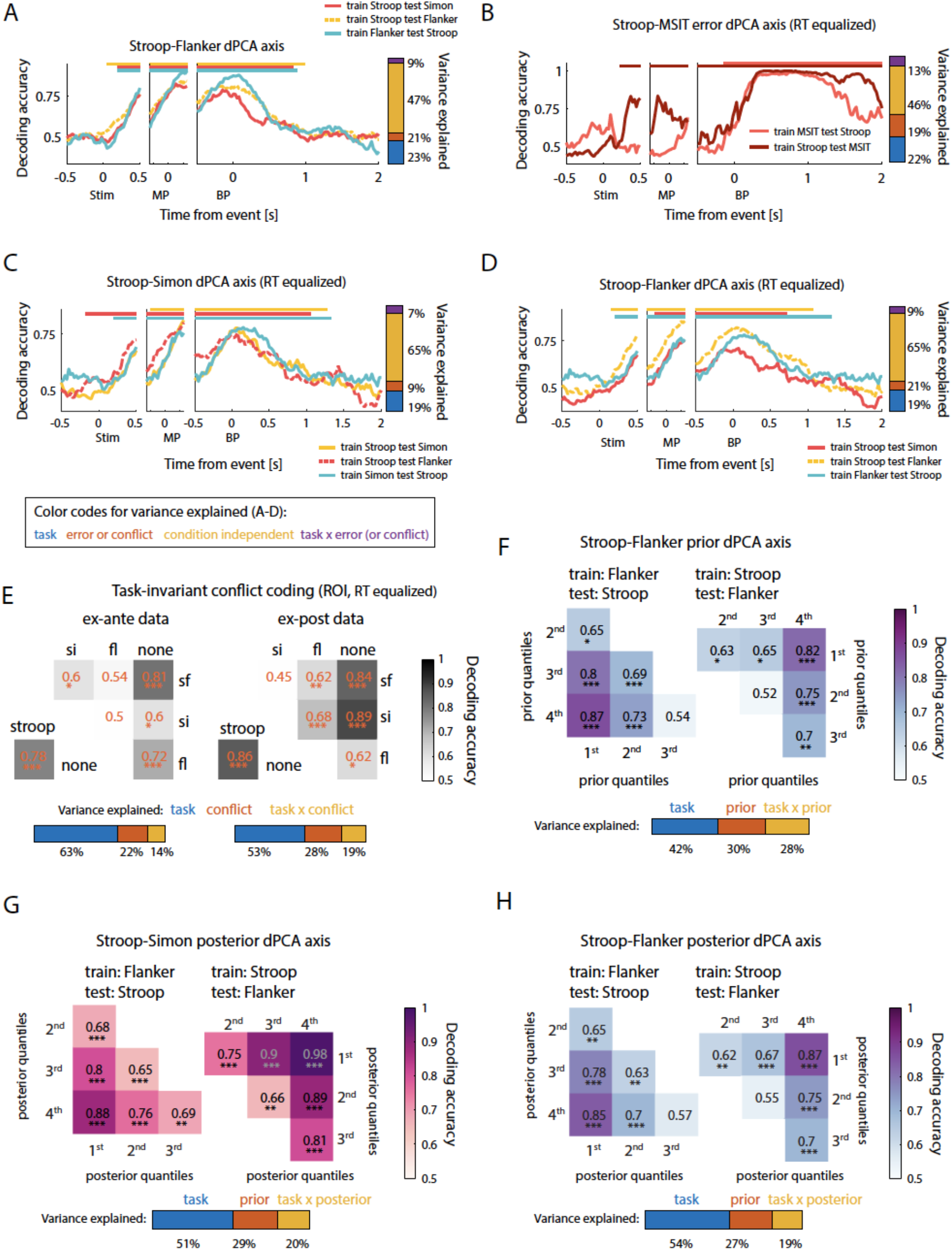
Task-invariant representation of performance monitoring signals. Because MSIT has two conflict conditions, Simon and Flanker, task invariance was investigated using Stroop/Simon and Stroop/Flanker separately. (A) Task-invariant decoding of conflict in MSIT (Flanker, yellow; Flanker red) and Stroop (green). (B) Task-invariant decoding of errors in MSIT (salmon) and Stroop (crimson) after RTs were equalized across conditions. The task-invariant coding dimension is extracted using dPCA that marginalizes out task information and time. This dPCA coding dimension is extracted from the error contrast in Stroop (error conflict vs. correct conflict trials) and the error contrast in MSIT (error “sf” trials and correct “sf” trials). Left, Accuracy for decoding errors as a function of time. Bar on the right shows the variance explained by the different dPCA components (color code see figure legend). (C-D) Task-invariant decoding of conflict in MSIT (Simon, yellow; Flanker red) and Stroop (green) after RTs were equalized across conditions. The dPCA coding dimension is extracted from conflict and non-conflict trials in Stroop and either from Simon and non-Simon trials (C) or from Flanker and non-Flanker trials (D), by marginalizing out task information and time. Left-out conflict trials and non-conflict trials in Stroop, and left-out Simon, non-Simon, Flanker, non-Flanker trials in MSIT are projected and classified by this coding dimension. Left, decoding accuracy of conflict as a function of time. The bar on the right shows variance explained by the different dPCA components (color code see figure legend). (E) Testing separability of conflict conditions in Stroop and MSIT using data from the ex-ante (left) and ex-post epochs (right) after RTs were equalize across conditions. The dPCA coding dimension used in this analysis is extracted by using conflict and non-conflict trials in Stroop and sf and non-conflict trials in MSIT by marginalizing out task information. Because data from ROIs are used, temporal information is already marginalized out before being used by the dPCA algorithm. Color coding represents decoding accuracy, orange numbers indicate the numerical values of decoding accuracy of that pair of conflict conditions. (F) Task-invariant decoding of all pairs of conflict prior levels in Stroop (lower triangle matrix) and Flanker (upper triangle matrix). The dPCA coding dimension here is extracted using the Stroop conflict prior contrast (the 1^st^ vs. 4^th^ quartile bins of Stroop conflict prior) and the Flanker conflict prior contrast (the 1^st^ vs. 4^th^ quartile bins of Simon conflict prior), marginalizing out task information. Color code represents decoding accuracy. Bar at the bottom shows variance explained of dPCA components. (G) Same as in (F) but for Stroop-Simon posterior. (H) Same as in (G) but for Stroop-Flanker posterior.

**Figure S10.**
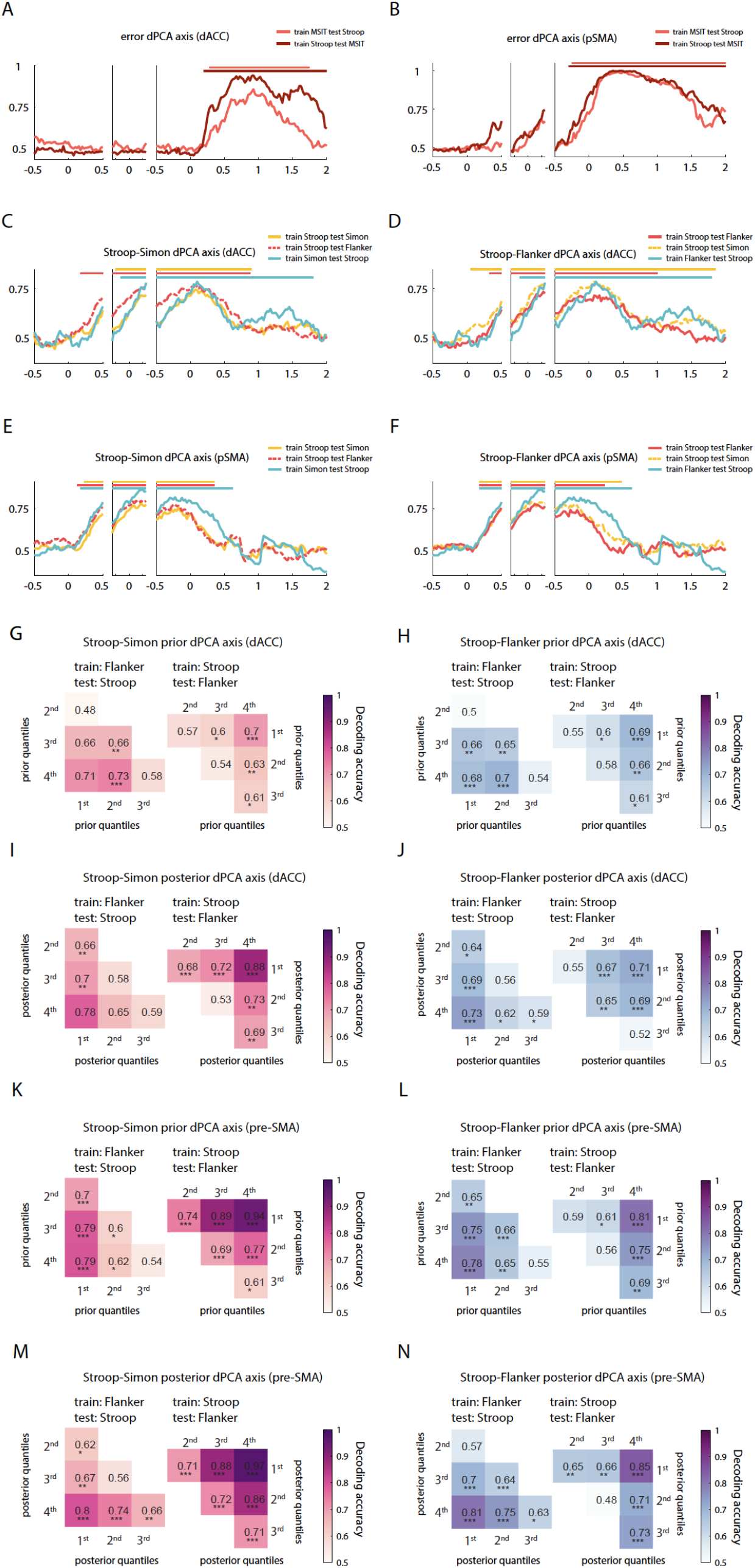
Domain-general representation of performance monitoring signals (separated for the dACC and the pre-SMA). (A) Task-invariant decoding of errors in both MSIT (salmon) and Stroop (crimson) using dACC neurons. This dPCA coding dimension is extracted from the error contrast in Stroop (error conflict vs. correct conflict trials) and the error contrast in MSIT (error “sf” trials and correct “sf” trials). This controls for trial conflict and isolates effects related only to error. Left, Accuracy for decoding errors as a function of time. Bar on the right shows the variance explained by the different dPCA components (color code see figure legend). (B) Same as in (A) but for the pre-SMA. (C-D) Task-invariant decoding of conflict in both MSIT (Simon, yellow; Flanker red) and Stroop (green), using dACC data. Because MSIT has two conflict conditions, Simon and Flanker, task invariance was investigated using Stroop/Simon and Stroop/Flanker conflicts separately. This dPCA coding dimension is extracted from conflict and non-conflict trials in Stroop and either from Simon and non-Simon trials (C) or from Flanker and non-Flanker trials (D), by marginalizing out task information and time. Left, decoding accuracy of conflict as a function of time. The bar on the right shows variance explained by the different dPCA components (color code see figure legend). (E-F) Same as in (C-D) but for pre-SMA data. (G-H) Task-invariant decoding of all pairs of conflict prior levels. The dPCA coding dimension here is extracted using the Stroop conflict prior contrast (the 1^st^ vs. 4^th^ levels of Stroop conflict posterior) and the Simon conflict prior contrast (the 1^st^ vs. 4^th^ quartiles of Simon conflict prior) in (G), and Flanker conflict prior contrast (the 1^st^ vs. 4^th^ quartiles of Flanker conflict prior) in (H), marginalizing out task information. Color code represents decoding accuracy. Bar at the bottom shows variance explained of dPCA components. (I-J) Same as in (G-H) but for conflict posterior. (K-L) Same as in (G-H) but for pre-SMA data. (M-N) Same as in (I-J) but for pre-SMA data. Horizontal bars demarcate the extent of significant clusters in time as determined by the cluster-based permutation test (p < 0.5).

**Figure S11.**
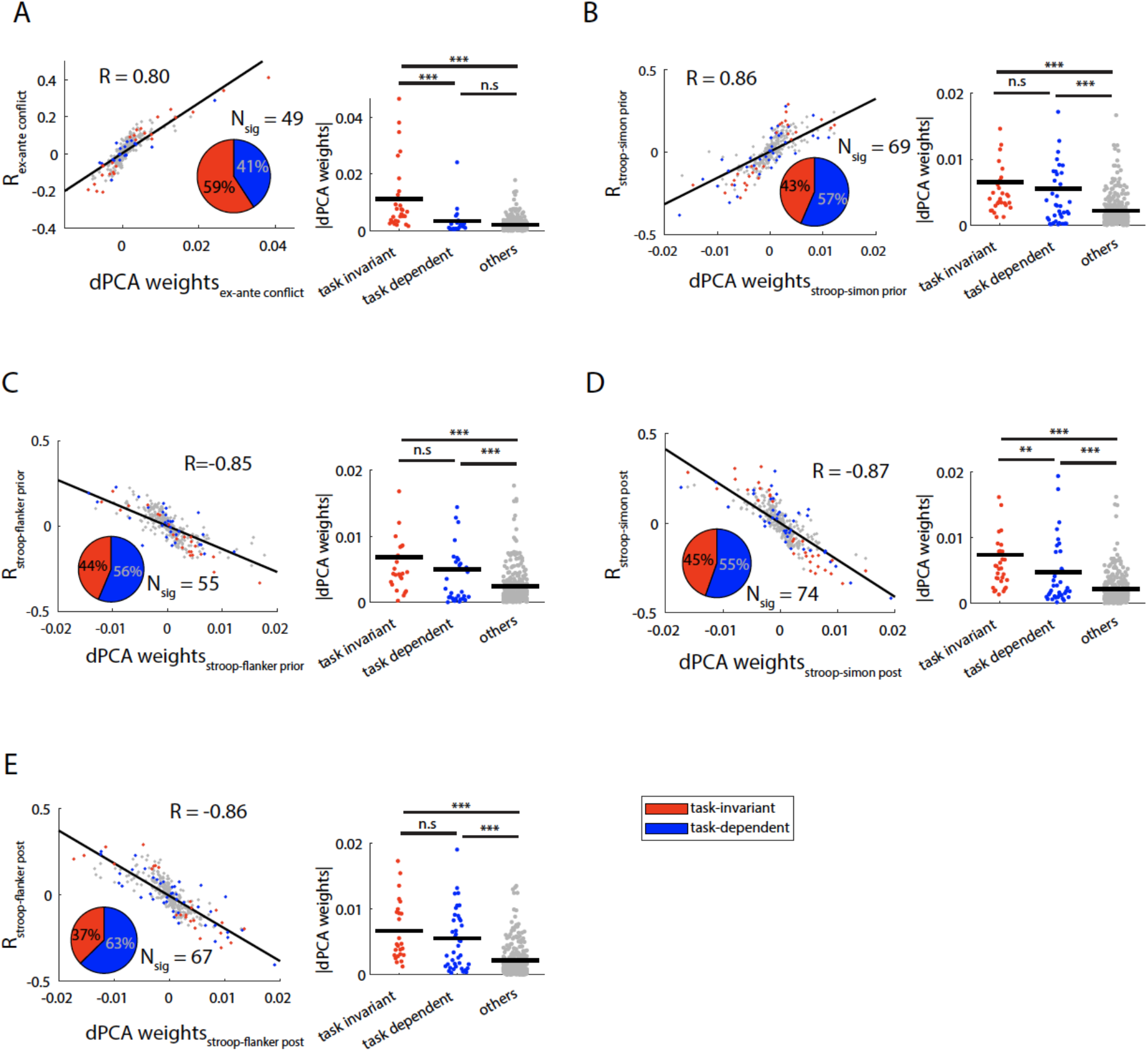
Relation between single-neuron tuning properties and dPCA weights. (A) Single-neuron contribution to task-invariant coding of conflict. Single-neuron coding strength of conflict is computed in the ex-ante epoch. (B-E) Relationship between single neuron tuning strength of Stroop-Simon prior (B) Stroop-Flanker prior (C), Stroop-Simon posterior (D), Stroop-Flanker posterior (E) and dPCA weights. The data used for computing single-neuron coding strength is the same as used in constructing the corresponding dPCA coding dimensions. Pie charts show the percentages of *tuned* neurons (total number is Nsig) that had a significant main effect for performance monitoring variable *only* (“task-invariant”, red slice) and those that had a significant interaction effect with the task identity (“task-dependent”, blue slice). Scatter plots (left) showed significant correlation between task-invariant coding strength and the corresponding dPCA weights. Right panel shows the regression weights between single neuron effect sizes and dPCA weights. *p < 0.05, ** p < 0.01, *** p < 0.001, n.s., not significant (p > 0.05).

**Table S1.**
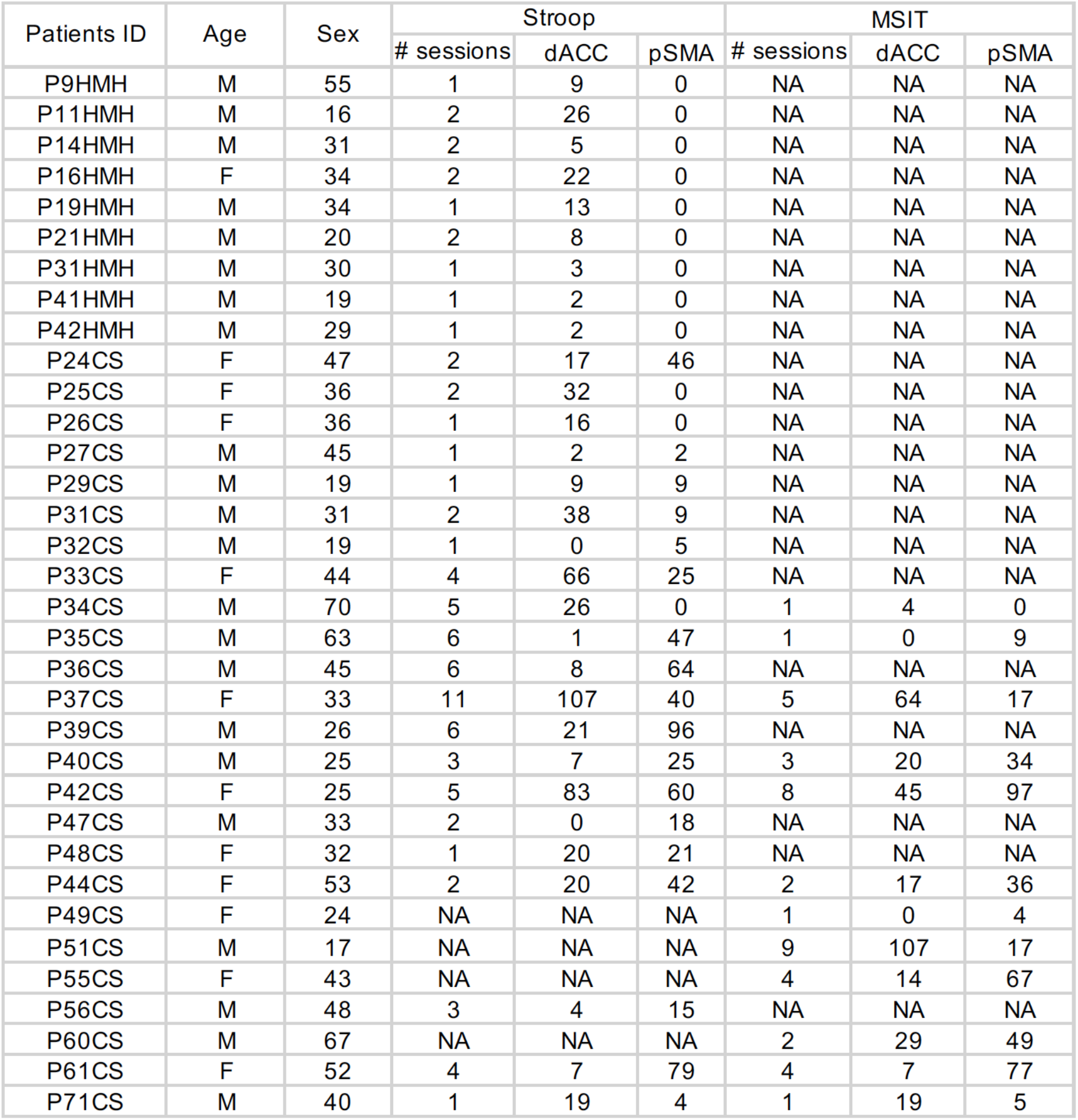
Number of sessions and neurons recorded. Summary of number of neurons recorded in each subject. For some subjects, both the Stroop task and MSIT were performed.

**Table S2.** Model comparisons for RT. Model comparison for RT using BIC. To test whether our RT-tuned Bayesian model explains variance in RT better than other models, we used linear mixed-effect models that considers subject variability (details of the model see **Methods**) and computed BIC for these models. The conflict prior is entered as a main fixed-effect and also as a by-session random effect. Here, conflict priors generated by four models are considered: “RT tuned”, Bayesian conflict learning model with DDM hyperparameters and thus the conflict prior is tuned by RT. “No RT tuning”, Bayesian conflict learning model without incorporating DDM likelihood for RT. “RL”, a reinforcement learning model where the conflict probability is modelled as a “value” function and updated trial-by-trial by a simple update rule. “Prev conflict”, a dummy variable indicating previous trial conflict. These linear mixed-effect models all have the same number of free parameters. A separate comparison was done between the RT tuned model with the model that uses the previous conflict (sub-table on the right) because the number of trials must be kept the same for the comparison and the “prev conflict” model did not consider the first trial for each session (there was no “prev conflict” in that case).

| BIC of RT |  |  |  |  |  |  |
| --- | --- | --- | --- | --- | --- | --- |
| Compare RT tuned with no RT tuning or RL |  |  |  | Compare RT tuned with previous conflict |  |  |
|  | RT Tuned | no RT tuning | RL |  | RT Tuned | Prev conflict |
| MSIT | -686.3 | -337.8 | -450.9 | MSIT | -805.7 | -440.1 |
| Stroop | -1412.1 | -904.1 | -972.5 | Stroop | -1733.6 | -1435.1 |

**Table S3.**
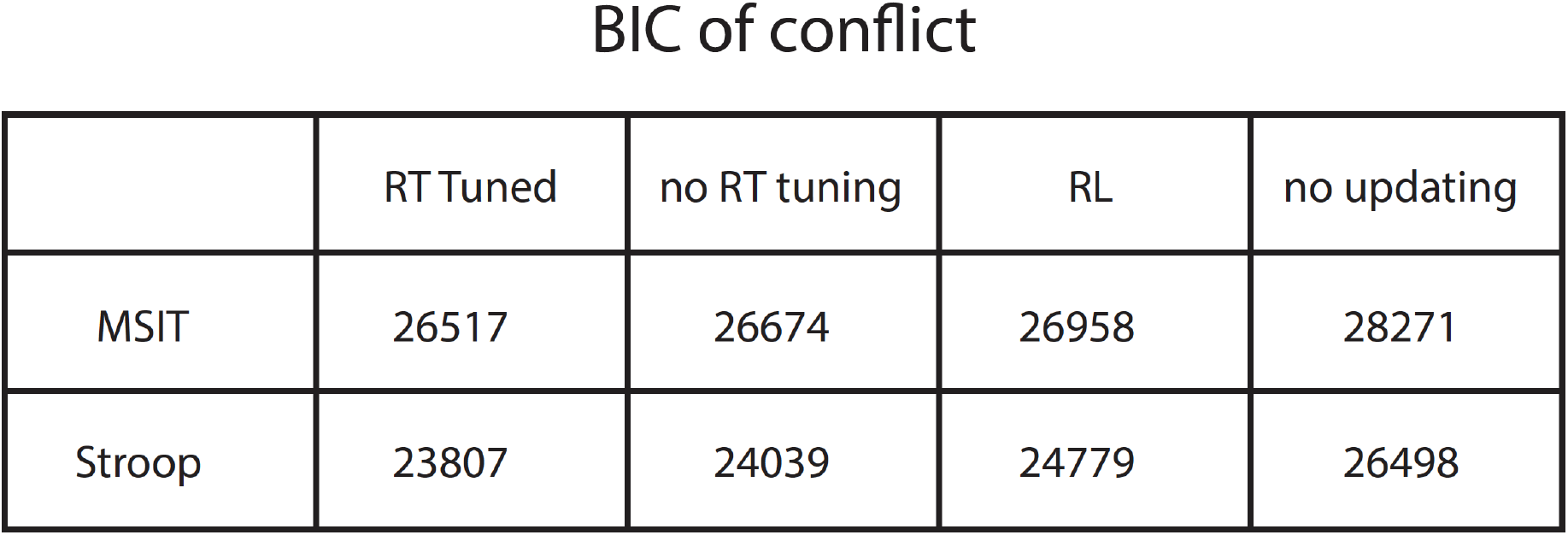
Model comparison for trial congruency. Model comparison for trial congruency using BIC. We used Bernoulli likelihood when computing BIC for the conflict sequence. Note that the number of fitted parameters for Bayesian models is zero, for the “RL” model is one (learning rate), and for “constant prior” model is one (the constant prior). BIC penalizes free parameters.

## Methods

### 1. Tasks

Subjects performed a speeded version of the Stroop and Multi-Source Interference (MSIT) tasks. For the Stroop task, subjects were shown a series of randomly intermixed color words (“red”, “green”, “blue”) printed in either red, green, or blue color (Fig. 1A). Subjects were instructed to name the color the word stimulus was printed in while ignoring its meaning and to do so as quickly and accurately as possible. For the MSIT task, subjects were shown an array of three numbers (0,1,2,3), out of which two were the same and the third of which was different (target). Subjects were instructed to press the button identical to the target number (which was unique) regardless of the position at which it was shown. For both tasks, all responses were recorded as button presses using an external response box (RB-740, Cedrus Corp., San Pedro, CA). For both tasks, the stimulus disappeared immediately when a button was pressed and was followed by a blank screen for 1s, followed by a feedback screen, which was shown for 1s. Subject were given three types of feedback: correct, incorrect or “too slow”. The feedback in 10-15% of trials (regardless of whether they were correct or incorrect) was “too slow” based on an adaptive response threshold (see (*34*) for details), which we used to emphasize the need to respond quickly and thereby resulting in a sufficiently large error rate. The inter-trial interval was sampled randomly from a uniform distribution between 1.5s to 2s. Trial sequences were pseudo-randomized and designed to avoid back-to-back repetitions of the same stimulus. The proportion of conflict trials in the Stroop task was 33%; For MSIT, the proportions of Simon only (“si”), Flanker only (“fl”), Simon and Flanker coincident (“sf”) trials are 15%, 15%, and 30%, respectively (the remaining 40% of trials have no conflict). The tasks were implemented in MATLAB (The Mathworks, Inc., Natick, MA) using Psychtoolbox-3 (*78*). The two tasks were performed in sequence, i.e., subjects finished blocks of one task first and then moved on to blocks of the other task. The order of task performed was randomized across experimental sessions.

### 2. Behavioral controls

As a control, we additionally collected behavioral data from N = 51 normal control subjects (24 females; age mean±sd: 44±11) using the Amazon mTurk platform. We implemented the MSIT task as described above using the jsPsych toolbox (*79*). These behavioral data were analyzed using the same methods as documented below. These control subjects exhibited a robust conflict prior effect like the patients (Fig. S1D).

### 3. Subjects

34 patients (see Table S1 for age and gender) who were evaluated for possible surgical treatment of their focal epilepsy using implantation of depth electrodes volunteered for the study and gave written informed consent. We only included patients with well-isolated single-neuron activity on at least one electrode in the dACC or preSMA. All research protocols were approved by the institutional review boards of Cedars-Sinai Medical Center, Huntington Memorial Hospital, and California Institute of Technology.

### 4. Electrophysiological recordings

We analyzed data from up to 4 electrodes in each subject (bilateral dACC and pre-SMA. For each depth electrode, there are eight microwires with high impedance microwires at the tip, and eight macro contacts with low impedance along the shaft (AdTech Medical Inc.). Data from all microwires and the most medial macro contact (which is placed within dACC or pre-SMA) are analyzed in this paper. For recordings from microwires, the sampling rate was 32-40kHz and the raw signal was acquired broadband (0.01Hz-9kHz). One microwire on each depth electrode was designated as a local reference wire. For intracranial EEG recordings done with macro contacts, the sampling rate was 2kHz (ATLAS, Neuralynx, Inc., Bozman, MT).

#### Electrode localization

Electrodes were localized using a combination of a pre-operative MRI and a postoperative MRI/CT using standard procedures we described elsewhere (*34, 80*).

Only electrodes that could be clearly localized to the dACC (cingulate gyrus or cingulate sulcus; for patients with a paracingulate sulcus, electrodes were assigned to the dACC if they were within the paracingulate sulcus or superior cingulate gyrus) or the pre-SMA (superior frontal gyrus) were included.

#### Spike detection and sorting

We filtered the raw broadband signal with a zero-phase lag filter in the 300-3000Hz band. Spikes were detected and sorted using a template-matching algorithm (*81*). Sorting quality is evaluated using the same metrics reported in (*34*) and only well-isolated single units are included in this paper. Channels with interictal epileptic events were excluded.

### 5. Quantification and statistical analyses

#### Behavioral modelling and analyses

We developed a Bayesian conflict learning model to infer subjects’ internal estimate of conflict probability (details see below). For this analysis, we concatenated all blocks of an experiments done in a single session. For trials with unusually long RTs (> 3 sd from the mean RT of the session), we replaced the outlier’s RT with the average RT computed from the neighboring 6 trials. We estimate the Bayesian model parameters using all trials but excluded error trials (after fitting) for analyses that focuses on conflict and conflict prior. We then analyzed whether the variance of RT was related to the estimated parameters using linear mixed-effect models (*82*). For MSIT, the linear mixed-effect model is specified as follows (all models are represented in Wilkinson’s notation):

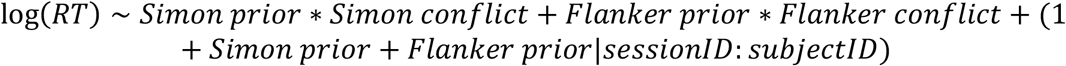

For Stroop, this is specified as:

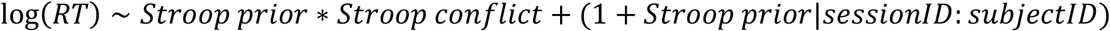

Here, the fixed effects of Simon, Flanker and Stroop conflicts are dummy variables indicating whether a particular trial involves conflict (value = 1) or not (value = 0). The fixed effects for priors are obtained from our Bayesian conflict learning models as detailed below. To test if RT was affected by conflict on the immediately preceding trial, represented by *Simon prevConflict, Flanker prevConflict* and *Stroop prevConflict,* we again constructed a linear mixed-effect models for both MSIT and Stroop. For MSIT, the model is specified as follows:

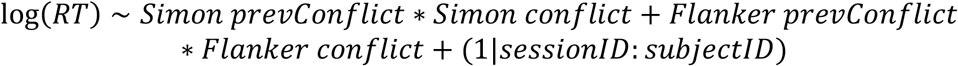

For Stroop, this is specified as:

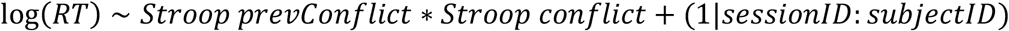

We investigated the effect of conflict prior on the likelihood of making an error using generalized linear mixed-effect models. For MSIT, this model is given as

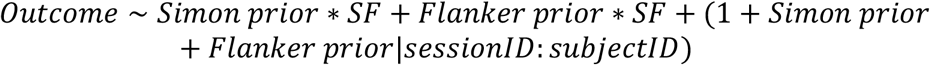

where *SF* is a dummy variable indicating whether the trial has both Simon and Flanker conflict (value = 1) or non-conflict (value = 0). We restricted this analysis to sf trials because most errors occurred on these trials. For Stroop, this model is given as

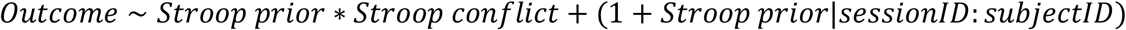

The response variable *Outcome* is a categorical variable indicating whether the trial ended in a correct (0) or incorrect (1) response.

To determine the statistical significance of each fixed effect, we compared the full model with a reduced model where the fixed effect in question was removed using the likelihood ratio test. To determine whether RT tuning of the model (see below) is necessary for conflict prior to explain RT variance, we replaced the conflict prior with the one estimated without RT tuning and kept all other terms the same. Statistical significance determined this way was indicated by stars or “n.s” (non-significant) in Figures 1 and S1. To determine whether the conflict prior explains variance in RT and error likelihood independent of practice, which is assumed to vary with the trial number, we augmented the aforementioned mixed-effect models by including three additional trial-ID terms: *trialID, trialID*^2^, *trialID*^3^ to capture variance related to the practice effects.

#### Bayesian conflict learning models

Our models are structurally similar to those used in several previous studies (*51, 54*). Here, we briefly highlight the modifications we made to extend these existing models to model behavior in both the Stroop and MSIT tasks, the latter of which has two types of conflicts that are monitored at the same time. Our models have the following parameters (see Fig. 1C for a schematic of the model structure): 1) a flexible learning rate *α*, which captures the subject’s belief in the rate of change in control demand in the environment (i.e., a change in conflict probability), and 2) conflict probability (*q_s_* for Stroop conflict in the model for Stroop, *q_si_* for Simon and *q_fl_* for Flanker conflicts in the model for MSIT). The models utilize two types of data (both of which are only available after a trial’s response has been made): 1) trial congruency *o* (value of 1 indicates an incongruent trial; *o_s_* for Stroop congruency, *o_si_* for Simon congruency and *o_fl_* for Flanker congruency; 2) reaction time *RT,* assigning Bernoulli likelihood function to the former and the drift-diffusion model (DDM) likelihood function to the latter (details see below). Prior work (*51*) uses a Gaussian likelihood function to describe RT generation, but we argue that the use of DDM has several advantages over the Gaussian approach: 1) fewer parameters are used in the DDM, making it computationally possible to model two types of MSIT conflict at the same time; 2) the DDM parameters have physiological meaning and thus also provide a clear physiological reasoning for conflict prior to affect specific components of the decision process; 3) DDM has been widely used and validated as the generative framework to model RT during decision making (*83, 84*).

Estimating control demand is operationalized as estimating the probability that a certain conflict (Stroop, Simon or Flanker) would occur in the block. One advantage of our models is that conflict probability and the rate of change in conflict probability are both estimated in an online manner, i.e., the models iteratively update their current estimates after every trial with new incoming data (of that trial) using Bayes’ law. This is consistent with the way human subjects perform conflict tasks while estimating the associated control demand: they perform and estimate trial-by-trial. Note that in this study the conflict probability was constant throughout the experiment (unbeknownst to subjects). Nevertheless, we allow the model to infer the learning rate *α* online because humans demonstrate inherent bias in believing that environmental statistics are not stable (*85*). There is therefore no fitting involved for estimating *α* and thus the models are not penalized for including this parameter in model comparisons. Note that we use a single *α* for both types of conflicts in MSIT. To simplify model estimation, we made the Markovian assumption that the current estimate of conflict probability depends only on the current trial congruency and RT, and the estimated conflict probability on the last trial, but not on the full history of past trial conflict probability (*54*). The iterative estimation of conflict probability then involves combining the estimated conflict probability from the previous trial (prior), transition functions capturing how the current estimate will change from the previous one (the probability of current estimate conditional on previous estimate) and the likelihood function.

The model starts with a transition function for the learning rate *α*:

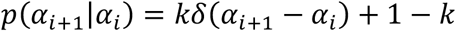

This formulation assumes that the learning rate has a probability k of having the same distribution as that of the preceding trial but with a probability 1-k of switching to a uniform distribution (over all possible *α*), because the learning rate is largely stable across time. The transition function for conflict probability concerning the transition from the current estimate to a future estimate is computed in two steps. Here, we refer to the current-trial estimate of conflict probability for Stroop, Simon or Flanker generically as *q_i_*, to which we assigned a uniform prior. The transition function is thus denoted as *p*(*q*_*i*+1_|*q_i_*, *α_i_*). First, an auxiliary variable *q*_*i*+0.5_ is constructed, which is a beta-distributed random variable with its mode being *q_i_* and the sum of two parameters being 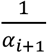:

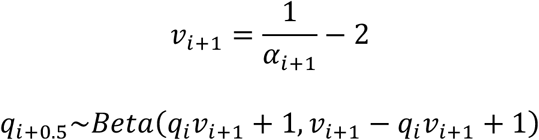

The conflict probability transition function is then constructed as

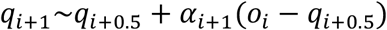

The transition function adopts a classical update rule used in reinforcement learning models, and the learning rate controls the balance between past *q*_*i*+0.5_) and current information (*o_i_*). For the MSIT model, we take the product of the transition functions computed separately for Simon and Flanker predicted conflict. Finally, we consider the likelihood function. Since the trial sequences were designed and re-used for different subjects, the estimated conflict probability would be the same across subjects for the same sequence, but this is inconsistent with the fact that such individual estimates should be subjective and different between participants. We thus incorporated RTs from each subject, which are assumed to be generated through a drift-diffusion process, to estimate a *subjective* conflict probability based on the assumption that a subjects’ RT reflects the extent to which they engaged control. RT is assumed to be generated by a diffusion process. The two bounds of DDM used here represent the correct and wrong choice (not response options). The diffusion process accumulated the *difference* in the evidence between the target and distractor response (Fig. 1C, right), which is smaller for conflict trials and thus leads to longer RTs. We refer to the drift-diffusion likelihood function (*86*) for RT as *p_DDM_* (*v_si_, v_fl_, v_sf_, v_non-conflict_, vz_si_, vz_fl_, a, q_si_, q_fl_*) for the MSIT model, and *p*_DDM_ (*v_confiict_, v_stroop non-conflict_, VZ, a, q_s_*) for the Stroop model. The hyperparameters specifying the DDM are boundary separation (*α*), base drift rates for Simon-only, Flanker-only, both Simon and Flanker present, and non-conflict trials in MSIT (*v_si_, v_fl_, v_sf_, v_non-conflict_*), base drift rates for conflict and non-conflict trials in Stroop (*v_conflict_, v_stroop non-conflict_*), and drift rate bias coefficients that scales the conflict probability of Simon, Flanker and Stroop (*VZ_si_, vz_fl_, vz*). The effective drift rate is then the sum between the base drift rate and the drift rate bias (Fig. 1C). Here we made the assumption that conflict prior affects RT by biasing *drift rates* based on a previous work investigating the effect of choice history on RT (*87*). We also assumed that the drift rate diffusion started at the half point of the boundary separation (i.e., *z* = 0.5). With the Markovian assumption, the updating process for the MSIT model is thus given by

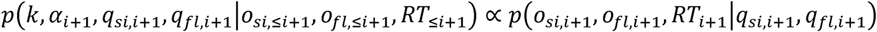

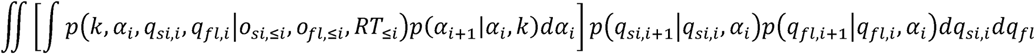

The updating process for the Stroop model is given by

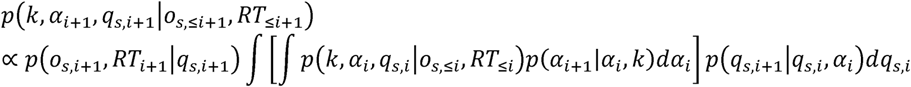

The likelihood function is the product of Bernoulli likelihood for trial congruency and DDM likelihood for RT. For MSIT, the likelihood function is given as:

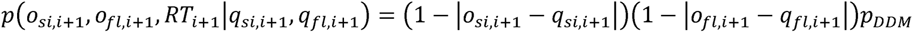

For Stroop, the likelihood function is given as:

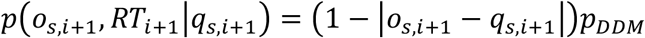

These hyperparameters are estimated using an expectation-maximization (EM) algorithm as shown in earlier work (*51*). Briefly, the model parameters were at first estimated *without* incorporating the DDM likelihood for RT (“E” step). Hyperparameters were then fit by maximizing the DDM likelihood for the observed RT using the conflict priors obtained (“M” step). The DDM likelihood function with the fitted hyperparameters were then incorporated into the Bayesian updating process (“E” step) to generate a new set of conflict priors, which were then used to maximize the DDM likelihood over observed RT again. These steps were repeated until the convergence of both model parameters and hyperparameters (Euclidean distance between parameter vectors from successive iterations < 10^-5^).

We considered three alternative classes of models: 1) reinforcement learning (RL) models; 2) constant model; 3) Bayesian learning model without RT tuning. For the RL model, we constructed a value function corresponding to the estimated conflict probability, and this estimate is also updated trial-by-trial using a Rescorla-Wagner rule. For MSIT, the update rule is:

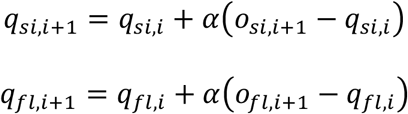

For Stroop, the update rule is:

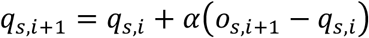

The free parameter *α* in the RL models was fit by maximizing the data likelihood (Bernoulli) for trial congruency. For the constant model, *q_s_, q_si_, q_fl_* were fit directly by maximizing the data likelihood for trial congruency. For Bayesian conflict learning models without RT tuning, *q_s_, q_si_, q_si_* were estimated online but the likelihood function for RT was not incorporated in the process.

We used the Bayesian Information Criterion (BIC) to compare the RT-tuned Bayesian conflict learning models with the RL models, constant models and the non-RT tuned Bayesian conflict learning models. We compared these models separately for their ability to explain trial congruency and RT. For this analysis, we pooled all data from all sessions and computed the BIC for each model and for each data type (RT or trial congruency), consistent with a previous study (*54*). Results of model comparisons can be found in Tables S2 and S3.

#### Selection of neurons

We defined the epochs of interests according to events in the tasks (Fig. 3A). The baseline epoch starts at 1.5s before stimulus onset and ends at stimulus onset. This epoch is used to analyze encoding of conflict prior. The ex-ante epoch is anchored to the midpoint of a period of time starting at 100ms after stimulus onset (to account for the minimal delay needed for visual information to reach the MFC) and ending at the time of button presses. We then defined the ex-ante epoch as a 500ms window centered on the midpoint of this period. The rationale for analyzing conflict signals in this epoch is as follows: at the early stage of stimulus processing, information about the different response options is not yet fully processed and hence minimal conflict; the conflict signal should reach its maximum when the different stimulus dimensions that drive competing responses are fully available; and finally, it should subside after a response is selected. We counted the number of spikes in these epochs and regressed the spike counts against the different regressors (error, conflict, conflict surprise, conflict prior and conflict posterior) using linear regression. For each regressor, we extracted a p value computed from the F test. A neuron was deemed selective for this regressor when p < 0.05. For MSIT, since there were both Simon and Flanker conflicts, neurons were selected when regressors related to either Simon or Flanker conflict were significant (e.g., ex-ante conflict cells in MSIT were the union of neurons selective for Simon conflict and Flanker conflict during the ex-ante epoch). To assess whether a neuronal class is significantly present in the population, we derive a null distribution by permuting the relation between spike counts and the regressor of interest for 1000 times. A p value is computed by comparing the true proportion of selected neurons against this null distribution. The 95^th^ interval of the null distribution for each neuronal class is plotted as dotted lines in Fig. 3B and Fig. S3A-B.

To statistically compare the extent of multiplexing between two groups of cells active in different epochs (Fig. 3D), we used the chi-squared test and reported the p-value and effect size of the test.

#### Single-trial spike train latency

We estimated the onset latency in individual trials using Poisson spike-train analysis (Fig. 3E) (*88*). This method detects the moments when the observed inter-spike intervals (ISI) deviate significantly from that assumed by a constant-rate baseline Poisson process. We used the spike rate averaged across the whole block of experiment as a baseline spike rates for each neuron. This baseline rate was then used to compute a Poisson surprise metric across the spike train. We started our detection algorithm from the onset of stimulus for each trial. For the ex-ante conflict neurons (two columns on the left), we restricted the range in which the detection algorithm looks for bursts to after stimulus onset and before button presses. This is because, by their definition, ex-ante conflict neurons carried a conflict signal before action. For the ex-post conflict neurons (two columns on the right), we restricted the range to 200ms before button presses and before end of trial. We then extract the latency of the first significant burst. The statistical threshold for detecting an onset was p < 0.01. Repeating the same procedure with a threshold of p < 0.001 did not affect our conclusions. For these analyses, we only used the conflict trials as we focused on the single-trial conflict response of selected conflict-encoding neurons.

#### Detrended fluctuation analysis

Detrended fluctuation analysis (DFA) was first developed by Peng and colleagues (*89, 90*) to quantify self-similarity in time series data. We use DFA to quantify the self-similarity in baseline spike counts on the scale of *trials.* First, the cumulative sum of the spike counts during baseline was computed. To be consistent with prior literature we refer to this cumulative sum as the *signal profile.* A set of trial window sizes were defined between the lower bound of 4 trials and the upper bound of the block length. For each window size, we then partitioned the signal profile into a series of data snippets. Partitioning was done such that two adjacent snippets had an overlap of half the window size. We then removed the linear trend from each data snipped (using least square regression) and computed the standard deviation across time. The mean of the standard deviations across all snippets of identical window size was then computed (y axis of Fig. S3H). Finally, the mean standard deviations were regressed linearly against the logarithmically scaled time windows and the slope was extracted as the DFA a value (Fig. S3H shows the fluctuations as a function of logarithmically transformed trial window sizes for two example neurons).

For Fig S3C-D, we tested the relation between a neuron’s baseline DFA a value and its tendency to encode conflict prior. To avoid selection bias, we split trials into two sets of equal size, with one half consisting of a consecutive run of trials. This is because DFA is used for time series data and thus required the data be consecutive and temporally ordered. For randomization, we first randomly sampled one trial from the first half of the block. Then a consecutive run of trials starting with this randomly picked trial as the starting point were extracted. The consecutive set was used to compute DFA a value while the rest of the trials were used to correlate with conflict prior (Simon, Flanker or Stroop) using Spearman rank correlation.

#### Decoding analysis (Support-vector machine)(Figs. S5,S6)

Data were aggregated from different experimental sessions to create pseudo-population data matrices. We constructed for each trial a peri-stimulus time histogram (PSTH) using 500ms bins in steps of 25ms. For all conflict or conflict prior-related decoding, we used correct trials only. Since different behavioral sessions had different number of trials (some subjects participated in less sessions than the others), we subsampled the same number of trials from each condition for each neuron and repeat this process 50 times.

For error decoding (Fig. S5A-B), we subsampled 10 error trials and 10 correct conflict trials for Stroop and 10 correct sf trials for MSIT. Since most errors occurred on high conflict trials, these contrasts isolate the effect of error while controlling for the effect of conflict.

For conflict decoding (Fig. S5C-E), we subsampled 30 trials from each conflict conditions: conflict and non-conflict trials for Stroop; Simon and non-Simon trials, Flanker and non-Flanker trials for MSIT.

For conflict prior (or posterior, Fig. S6), trials were binned by quartiles of conflict prior (or posterior) into four bins (“levels”) separately. For each pair of levels, one decoder was trained and tested. This analysis used spike counts from the ex-ante or ex-post epochs (ROIs).

For each time bin or ROI, we performed 5-fold cross validation using LIBSVM (*91*). We used the linear kernel and set the *c* parameter to 1 for all analyses. In brief, trials were first randomly split into 5 equal parts; each part was used in turn as the testing data while the rest of the four parts were used as training data. Decoding accuracy was the proportion of correct classifications among the 250 samples (50 resamples x 5 folds). Note that the resampling was done once to generate testing and training sets for the whole time series and used for both within-time and across-time decoding. For within-time decoding (All plots with dotted lines in Fig. S5 and plots within dotted frames in Fig. S6), the SVM classifier trained using the training data from each time bin was tested using the testing data from the same time bin. For cross-temporal decoding (temporal generalization; all plots with solid lines in Fig. S5 and plots without dotted frames in Fig. S6), the SVM classifier trained using the training data from each time bin were tested across the trial using testing data gathered from other time bins.

#### Reaction times equalization

For analyses shown in Fig. S9B-E, we selected a subset of trials from each condition so that the RTs did not differ significantly across conditions (e.g., equalizing RTs between conflict and non-conflict trials in the Stroop data). Here we detail the RT equalization procedure we used to create “RT equalized sets”. We first selected a condition as the “anchor” condition. We sorted the RTs of the anchor condition, and for each RT we searched in the target (to-be-equalized) condition(s) for a trial whose RT did not differ from the anchor RT by more than 0.1s. If all RTs in the target condition differed from the anchor RT by > 0.1s, the anchor RT was not included in the RT equalized set. Once selected, the anchor RT and the target RT were both removed from the original set to ensure that no trials were included twice in any RT equalized trial sets. This procedure was repeated until one of the conditions considered were emptied. We confirmed post hoc that RT equalization was successful by testing whether RTs were not significantly different using ANOVA (p > 0.5 for all the RT equalized sets).

#### Decoding analysis (population activity vectors and demixed PCA)

Data were aggregated from different experimental sessions to create a pseudo-population. We randomly selected one trial for each neuron from one condition and concatenate the data from each neuron to form a single-trial testing data matrix. The rest of the trials were averaged for each condition and concatenated to form a training data matrix. Coding dimensions were defined based on the condition-averaged training data. To define the coding dimensions used to decode conflict conditions within MSIT, we used the population activity vectors (a high dimensional vector in the raw firing rate space) defined by the difference between the two condition means. To define coding dimensions for the cross-task decoding problems we used dPCA to extract demixed principal components (dPC). Details of which trials were used to define the coding dimensions are given in the sections to follow. Both testing and training data were projected onto the identified coding dimensions. The labels for testing data were assigned according to the label of the nearest neighbor of the training data. To test condition generalization, we projected the testing data from one pair of conditions to a coding dimension defined by another pair of conditions (e.g., a Simon trial and non-Simon trial projected to the population vector flanked by Flanker and non-Flanker trial averages) and classified using the labels of the nearest projected training data. This decoding procedure was repeated 1000 times (resulting in 1000 single-trial testing data matrices and the corresponding training data matrices), and the decoding accuracy was defined by the proportion of correct classifications among these 1000 repetitions. To determine statistical significance, we permuted the trial labels for 500 times and for each permutation, we repeated all above steps to generate a null distribution. A p-value was computed from comparing the true decoding accuracy with the null distribution.

#### Pseudo-population matrices for MSIT state space analyses

For Fig. 4E-G, trials were binned by quartiles of conflict prior (or posterior) into four bins (“levels”) separately. However, because conflict prior was updated into conflict posterior after each button press, binning priors does not guarantee that the posteriors would fall into the same bins. This is because updating is specific to each behavioral session and thus differs between neurons. Averaging trials using only bins formed by prior quartiles would thus mix trials with different levels of posterior for each neuron. To avoid this problem, we thus formed the data matrix (which now includes the time dimension rather than a single ROI; spikes were counted in 500ms bins swept across the whole trial in steps of 25ms; spike trains were aligned to button presses) by concatenating two submatrices: one that was constructed by averaging trials within bins defined by prior quartiles using data *before* button presses, and one that was constructed from averaging trials within bins defined by posterior quartiles for neural data *after* the button press. We then used PCA to find the three PCs that explained the most variance for this matrix. The concatenated data matrix was then projected onto these PCs to generate the visualization of trajectory corresponding to prior (or posterior) levels.

#### Population coding dimensions for conflicts within MSIT

We next describe how coding dimensions were defined in each case using population activity vectors in the raw firing rate space. For Fig. 4B, the coding dimension was the population vector flanked by the trial averages of sf and non-conflict trials (Fig. 4A, dashed lines). Classifications were carried out between pairs of conflict conditions (e.g., between si and fl trials) as detailed above. For Fig 4C, we took a bin-wise approach to investigate whether Simon conflict representation generalize to Flanker representation, and vice versa. For this, we split trials into four non-overlapping groups: Simon, Flanker, non-Simon, non-Flanker trial sets. We split sf and non-conflict trials randomly in half. One half of sf trials were pooled with si trials to form the Simon trial set, and one half of non-conflict trials were pooled with fl trials to form the non-Simon trial set. The other half of sf trials were then pooled with fl trials to form the Flanker trial set, and the other half of non-conflict trials were pooled with si trials to form non-Flanker trial set. Using these trial sets, for each time bin we extracted two coding dimensions from the training data: one population vector flanked by trial averages of Simon and non-Simon trials (Simon coding dimension), and one population vector flanked by the trial averages of Flanker and non-Flanker trials (Flanker coding dimension). We then projected the testing data from Simon and non-Simon trials onto the Flanker coding dimension and classified the testing data using the closest projected training data, and vice versa. For details of this classification procedure see above paragraph. This assesses the extent to which coding of Simon and Flanker conflict is abstract.

#### Compositionality of conflict representations within MSIT

For Fig. 4D, the coding dimensions were taken to be the blue and orange edges as shown in Figure 6A. The purpose of this analysis is to assess to what extent the representation of conflict is compositional (within a task). We assumed that in the neuronal firing rate space, the representation of Simon/Flanker conflict is a vector pointing from non-conflict trial averages to the si/fl trial averages. Compositionality of such conflict representation would imply that the sf representation (vector pointing from non-conflict trial average to the sf trial average) is equal to the sum of the Simon and Flanker representations. According to the parallelogram law of vector addition, this then corresponds to the blue and orange edges in Fig. 4A forming a parallelogram. We tested the extent of parallelism in the data using decoding. The coding dimensions here were defined by the following population vectors using training data: one flanked by non-conflict and si trial averages (Fig. 4A, blue), one flanked by fl and sf trial averages (blue), one flanked by fl and non-conflict trial averages (orange) and one flanked by si and sf trial averages (orange). Left-out testing data from conditions flanking one of the blue or orange pair of edges were then projected to the other edge in the pair and classified by the training data defining this edge. For example, single-trial testing data of non-conflict and si trials were projected to the coding dimension flanked by fl and sf trial averages and were classified by fl or sf trial averages.

#### Relationship between single neuron tuning and the compositional conflict representations within MSIT

For Fig. S8E-F, the goal is to investigate the relation between the nonlinearity in single neuron conflict coding and the deviation from compositionality in state space representation of conflict. We denote the state-space representation of Simon and Flanker - only conflict as the population vectors flanked by the trial averages of si and non-conflict and by the trial averages of fl and non-conflict. We refer to the state space location occupied by the linear sum of Simon and Flanker representation defined above as “s+f”. The deviation from perfect compositionality is then given by the population vector flanked by “sf” and “s+f”. The loading of “sf” to the “s+f” vector reflects the single neuron contribution to the deviation at the population level. To quantify nonlinearity of conflict coding for each neuron, we first regressed the spike counts in the ex-ante or ex-post epoch (1s) against three fixed effects: a Simon effect (dummy variable indicating the presence or absence of Simon conflict on a trial), a Flanker effect (dummy variable indicating the presence or absence of Flanker conflict on a trial) and the interaction term between these two. We extracted the F statistic related to the interaction term, which captures the effect of nonlinear mixing of Simon and Flanker conflict. We then extracted a population vector flanked by the sf trial average and and “s+f”, the sum of two population vectors one flanked by trial averages of si trials and non-conflict trials, and one flanked by trial averages of fl trials and non-conflict trials. We then correlated the loading of “sf” - “s+f” vector and the F statistics from a particular neuron.

#### Quantification of state space dynamics

For Fig. S8G-I, we binned spike counts using 250ms bins swept across the trial in steps of 10ms. The state-space speed was defined to be the Euclidean distance between population vectors of adjacent time bins divided by the step size. We averaged the state-space speed across time within an epoch. We also computed the Euclidean distance between pairs of trajectories (1^st^ and 2^nd^, 2^nd^ and 3^rd^,3^rd^ and 4^th^) and averaged this across trajectories and across time bins within an epoch. State-space speed and the averaged distance between trajectories were plotted against each other in Figure 6I. Our method for extracting speed in state-space follows prior work (*92*).

#### Testing ordinal relationship of prior (or posterior) projections

We analyzed the ordinal relation between neural projections of prior (or posterior) as shown in Fig. S8J-L. PCA axes encoding prior/posterior variance were extracted from spike count data collected in ROIs (baseline for prior and the ex-post epoch (0-1s after button presses) for posterior). Since prior/posterior is continuously valued, we created four trial conditions by binning the trials using quartiles of prior/posterior. For each type of prior or posterior (Simon, Flanker and Stroop), we projected the left-out trial (not used for computing the PCA axis) onto the PCA axis for each trial condition and this procedure was repeated 1000 times, yielding 1000 projected values for each trial condition. We then regressed the projected values (concatenated into a vector) against their trial condition labels (1^st^,2^nd^,3^rd^,4^th^ quartile bins) using a multinomial logistic regression with the assumption of ordinal relation between trial groups. Essentially, we were testing whether the out-of-sample project values can reliably predict the trial condition they belong to assuming that the conditions were ordinal. We reported the p-value and t-statistic of the effect of projected values.

#### Demixed Principal Component Analyses (dPCA)

We used dPCA to extract task-invariant representation of performance monitoring signals. We use “s”, “si” and “fl” to denote Stroop, Simon and Flanker conflict condition, respectively. We use “sf” to denote the condition where both Simon and Flanker conflict are present.

Analyses on conflict and conflict prior (or posterior) used only correct trials. We used dPCA as described previously (*56*), with the following adaptions. The dPCA algorithm first decomposes population neural activity into marginalized data matrices with respect to the variables of interest. We constructed the marginalized population activity (referred generically as 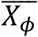) with respect to error (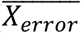 in Fig. 5A) or conflict (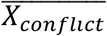, Fig. 5B, Fig. S9A) by marginalizing out *time* and *task identity* dimensions (denoted by “〈·〉_*task,t*_”).

For interpretability, we investigated whether the neural representation is abstract across tasks separately between Stroop and Simon conflict (“*s* & *si conflict*” is an indicator variable marking the presence or absence of conflict in either tasks) and between Stroop and Flanker (“*s* & *fl conflict*” is an indicator variable marking the presence or absence of conflict in either tasks). Set up this way, the “task” dimension captures variance related to task set differences (Stroop vs. MSIT). To compute marginalized averages, we use N-dimensional population activity

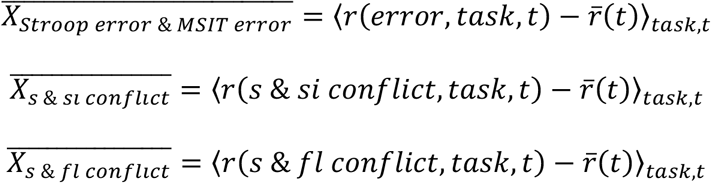
where 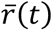 is the firing rate averaged across trials and time bins. For Figure S9B-D, these definitions are the same except that RT equalized trial sets were used.

For Figure S8A-B, we sought a common coding dimension between error and conflict separately in MSIT and Stroop, by marginalizing out the information about time and which pair of conditions were contrasted (denoted by “〈·〉_*contrast,t*_”; The “contrast” indicator, for MSIT it indicates whether the contrast considered is sf vs. non-conflict or error sf vs. correct sf; for Stroop it indicates whether the contrast considered is correct conflict vs. correct non-conflict or error vs. correct conflict). For Fig S8A-B, we constructed the marginalized population activity with respect to error vs conflict in both MSIT (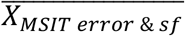, *MSIT error* & *sf* is an indicator variable marking the presence or absence of errors and MSIT sf conflict) and Stroop (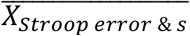, *Stroop error & conflict* is an indicator variable marking the presence or absence of errors and Stroop conflict) as follows:

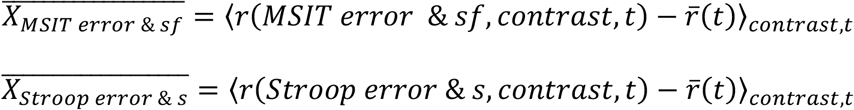

For analyses in Figs. 5C-D, S9F-H and S10G-N investigating task-invariant coding of conflict prior and conflict posterior, we used data from *a single ROI* (ex-ante or ex-epoch) and hence only the task but not time dimension was marginalized out. For Fig. 5C, we investigate task invariance between Stroop conflict and MSIT sf conflict (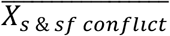, *s* & *sf conflict* is an indicator variable marking the presence or absence of Stroop conflict and MSIT sf conflict):

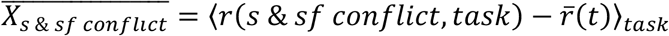

For analyses in Fig. S9E, this definition the same except that RT equalized trial sets were used.

Here again for interpretability, we investigated task invariance between Stroop and Simon prior (or posterior; Figs. 5D, S9G, S10G,I,K,M) and between Stroop and Flanker prior (or posterior; Fig. S9F,H, S10H,J,L,N) separately (e.g., 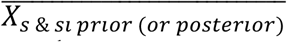, *s* & *si prior* (*or posterior*) is an dummy variable marking the 1^st^ and 4^th^ levels of Stroop and Simon prior (or posterior); 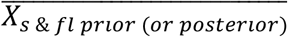, *s* & *fl prior* (*or posterior*) is an dummy variable marking the 1^st^ and 4^th^ levels of Stroop and Flanker prior (or posterior)):

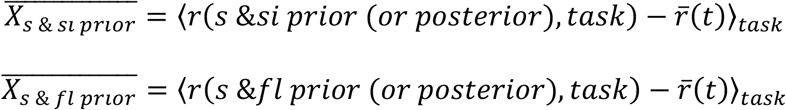

The algorithm then finds encoding (*F_ϕ_*) and decoding (*D_ϕ_*) matrices separately for each marginalized averages using the regularized reduced-rank regression:

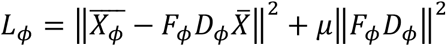

We assigned a fixed regularization coefficient *μ* to avoid overfitting (*μ* = 6*e*^-6^ determined from results reported in (*56*)). We used the columns of *D_ϕ_* as the demixed principal components (dPC) and projected N-dimensional data (single-trial data for testing and trial-averaged data for training) to these dPCs. The numerical values of *D_ϕ_* reflects the contribution for each neuron to task-invariant representation.

To test the statistical significance of coding dimensions, we randomly chose one trial for each trial type (e.g., one error trial and one correct trial) and constructed a single-trial activity matrix *X_test_*. We then used the remaining trials to form the trial-averaged training data 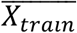, which is used to find the dPCA coding dimensions. The left-out single-trial data *X_test_* is then projected onto the first coding dimension that captures the most variance computed from *X_train_*, and classified according to the closest class mean. We repeated this procedure 1000 times and determined the decoding accuracy as the proportion of correct classification among the 1000 test trials. We then generated the null distribution by shuffling the trial labels and then repeated the decoding procedure 500 times. For Figs. 5C-D, S9E-H, statistical significance is determined by comparing the true decoding accuracy with this null distribution. For Figs. 5A-B, S8A-B, S9A-D, S10A-F, statistical significance is determined by the cluster-based permutation test using this null distribution (*93*). The fraction of explained variance (Bars in Figs. 5A-D, S9A-H) for each marginalization is given by:

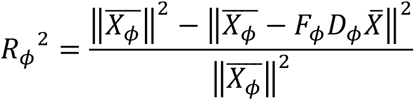

#### Single-neuron contribution to task-invariant coding dimensions

For Figs. 5E-F and S11, we quantified classified neurons by the extent of task invariance in their coding profile. Spike counts from two tasks were aggregated and regressed against a task identity dummy variable (0: Stroop, 1: MSIT), performance monitoring variable, and the interaction between these two regressors. Neurons with significant performance monitoring variable, but insignificant task identity were named “task-invariant”. Neurons with significant interaction term were named “task-dependent”. Neurons without any significant terms were named “others”. We quantified the signed single-neuron coding strength by a partial correlation (used in the scatter plots in Figs. 5E-F and S11): correlation between spike counts and performance monitoring variable, after partialing out the task identity.

Performance monitoring variables can be error (error in either task is marked 1 and correct trials in either task is marked 0), conflict (conflict in either task is marked 1 and non-conflict in either task is marked 0), conflict prior (or posterior; 1^st^ level in either task is marked 0 and 4^th^ level in either task is marked 0).

